# Horizontal transfers between fungal *Fusarium* species contributed to successive outbreaks of coffee wilt disease

**DOI:** 10.1101/2023.12.22.572981

**Authors:** Lily D. Peck, Theo Llewellyn, Bastien Bennetot, Samuel O’Donnell, Reuben W. Nowell, Matthew J. Ryan, Julie Flood, Ricardo C. Rodŕıguez de la Vega, Jeanne Ropars, Tatiana Giraud, Pietro D. Spanu, Timothy G. Barraclough

## Abstract

Outbreaks of fungal disease have devastated plants and animals throughout history. Over the past century, the repeated emergence of coffee wilt disease caused by the fungal pathogen *Fusarium xylarioides* severely impacted coffee production across sub-Saharan Africa. To improve the disease management of such pathogens, it is crucial to understand their genetic structure and evolutionary potential. We compared the genomes of 13 historic strains spanning six decades and multiple disease outbreaks to investigate population structure and host specialisation. We found *F. xylarioides* comprises at least four distinct lineages: one host-specific to *Coffea arabica*, one to *C. canephora* var. *robusta*, and two historic lineages isolated from various *Coffea* species. Mapping variation onto a new long-read reference genome showed that host-specificity appears to be acquired through horizontal transfer of effector genes from members of the *F. oxysporum* species complex. This species complex is known to cause wilt disease in over 100 plant species. Multiple transfers into the *F. xylarioides* populations matched to different parts of the *F. oxysporum* mobile pathogenicity chromosome and were enriched in effector genes and transposons. Effector genes in this region and other horizontally transferred carbohydrate-active enzymes important in the breakdown of plant cell walls were shown by transcriptomics to be highly expressed during infection of *C. arabica* by the fungal arabica strains. Widespread sharing of specific transposons between *F. xylarioides* and *F. oxysporum*, and the presence of large *Starship* elements, indicate that transposons were involved in horizontal transfers. Our results support the hypothesis that horizontal gene transfers contributed to the repeated emergence of this fungal disease.

## Introduction

Fungal diseases have devastated plants and animals throughout history, with the first records dating from biblical plagues [1]. Many examples of famines and harvest failures result from outbreaks of fungal disease with wide-ranging and important human consequences, and modern farming practices risk inadvertently helping the emergence of new pathogens [2]. Coffee is a major commodity that is worth over $22 billion to the global economy [3], provides high export earnings (e.g., one third exports in Ethiopia [4]) and supports millions of smallholder farmers [5]. However, as with other crops, there is a constant threat from pests and disease, especially fungal pathogens. Coffee wilt and leaf rust have caused major losses to yields and livelihoods in Africa and central America respectively [6, 7]. Pathogens can rapidly evolve [2], so understanding how new variants emerge is vital to developing sustainable disease control measures.

Coffee wilt disease, caused by the fungal pathogen *Fusarium xylarioides*, infects coffee plants via roots or cuts to colonise the xylem [8]. Wilting results from the pathogen blocking water uptake, and cell walls are degraded for nutrients [9]. Currently, two host-specialised strain types are known to infect the main African coffee crops: arabica (*Coffea arabica*), grown at high altitudes; and robusta (a variety of *C. canephora*), grown at low-mid altitudes in tropical Africa [8]. Coffee wilt disease has a well-documented history of emergence and outbreaks since its discovery in 1927 [10]. By the 1950s, coffee crops in west and central Africa were decimated, resulting in the replacement of commercial cultivars with resistant robusta coffee [6]. Yet by the 1970s, coffee wilt disease re-emerged on robusta coffee in central Africa [11], becoming a major problem in east and central Africa by the mid-1990s. Production did not recover in some countries (e.g. Democratic Republic of Congo, [12]). In parallel, coffee wilt disease was first reported on arabica coffee in Ethiopia in the 1950s, becoming widespread in the 1970s, but with low severity of disease [13]. However, recently disease severity in Ethiopia has increased [14, 15]. Key questions for understanding the emergence of coffee wilt disease include: what are the genetic relationships among the strain types on different hosts and at different times; and what genetic changes underlie their ability to infect different hosts? New pathogen variants often bring together gene pools present in different genomes [16]. Could new effectors emerge from gene pools shared across a population on a given host, multiple populations on different hosts, or completely different species?

There is growing evidence that horizontally transferred regions (HTRs) between distantly related fungi are important for the emergence of new pathogen variants [17]. For example, a 14kb fragment encoding the ToxA virulence protein and abundant transposons was transferred between three wheat fungal pathogens of the order *Pleosporales* [18]. This fragment was later identified as a *Starship*, which are large mobile elements found across every major class of filamentous ascomycetes, moving large sets of genes within genomes, species, or even between species, and which can conflict or cooperate with their fungal host genome [19]. Another example is *F. oxysporum*, which causes wilt disease in over 100 plant species with host-specific *formae speciales* [20]. Mobile supernumerary chromosomes with “supercontigs”, namely sub-regions enriched in secreted in xylem *SIX* effector genes and *mimp* transposons, a type of miniature-inverted *impala* repeat, transfer between *formae speciales* and determine host-specificity [21–23]. Transposons are hypothesised to promote hyper-variability of sets of effector genes, either directly by transposition or by facilitating recombination to generate new gene assortments [24]. Thus, while fungal populations may generally evolve within local gene pools, horizontal transfer between more distant relatives can introduce diversity and enable outbreaks of new types of pathogen [25].

A previous study sequenced six *F. xylarioides* genomes comprising arabica, robusta and strains isolated in 1950s which we describe as “coffea” for their ability to infect multiple *Coffea* species [26]. Genome-wide variation supported an early split between arabica and other strain types, consistent with pre-agricultural divergence on wild coffee species, and potentially a separate gene pool on *C. arabica*. In contrast, the robusta strain type from the 1970s onwards appeared to derive from the pre-1970s coffea strain type outbreak. Different strain types featured different putative effector genes, whereby some effectors showed close matches with *F. oxysporum*, but were absent from closer relatives of *F. xylarioides*. Strikingly, a 20 kb scaffold showed high sequence similarity to an *F. oxysporum* pathogenicity chromosome and contained the same genes and *mimp* transposons, also lacking in closer *F. xylarioides* relatives. This led to the hypothesis that host specialisation and changes in subsequent outbreaks involved diverged strain types receiving HTRs from several *F. oxysporum* formae speciales and thus a greatly expanded gene pool of wilt-associated effector genes.

Here, we test this hypothesis using comparative genomics of eleven *Fusarium* genomes, including *F. oxysporum* and *F. solani* chosen to represent potential HTR donors in crops that grow with or near coffee in Africa (Table S1). Aligning *F. xylarioides* genomes with a new long-read reference genome for the arabica strain type, we found that the different outbreaks on diverse coffee types arose as separate populations. The populations differ in the presence and absence of large genomic regions with synteny and high sequence similarity with multiple different *F. oxysporum* ff. spp; this is consistent with horizontal acquisition from distantly related lineages. Transcriptomics revealed that putative effector genes are highly expressed during coffee plant infection, with several putatively originating from *F. oxysporum* and absent in *F. xylarioides* sister species. Finally, all *F. xylarioides* populations share highly similar *mimps* and other transposable elements associated with *F. oxysporum*’s mobile pathogenic chromosome, that could have mediated horizontal gene transfer (HGT) between them. Together, these results support the hypothesis that HGT contributed to the emergence of host-specialised populations that defined successive outbreaks of coffee wilt disease.

## Results

### Phylogenomic support for *Fusarium xylarioides* as a distinct, monophyletic species complex

Short-read genome sequencing supported the delimitation of *F. xylarioides* as a species complex. Analysing 11 new Illumina genome assemblies of five *F. xylarioides* strains (Table S1, Figure 1) and six public *Fusarium* genomes (Table 2), together with 25 published *Fusarium* genomes (Table S2), assigned protein-encoding genes to 26,000 orthologous groups, of which 3544 had members in all genomes with 1685 single-copy genes. A species tree based on these 3544 orthogroups supported phylogenetic relationships in the *Fusarium* genus from earlier studies with lower sampling [26, 27], Figure 2A). *Fusarium xylarioides* forms a monophyletic clade within the *F. fujikuroi* species complex (Figure 2A), a result that is supported by the majority of gene trees (94%) (Figure S1) and by the high average coding sequence identities among *F. xylarioides* (98%) contrasting with lower similarity with *F. phyllophilum* (96%), other species of the *F. fujikuroi* complex (91-92%) and *F. oxysporum* strains (90%) (Figure 2B). Multilocus species delimitation, based on congruence among gene trees, identified phylogenetic units that matched the named species except for one strain of *F. oxysporum*. Within the *F. xylarioides* species clade, there were four well-resolved groups: arabica, robusta, and two separate clades differentiating the coffea genomes (those isolated from the pre-1970s outbreak) (Figures 2 and S1).

**Figure 1:**
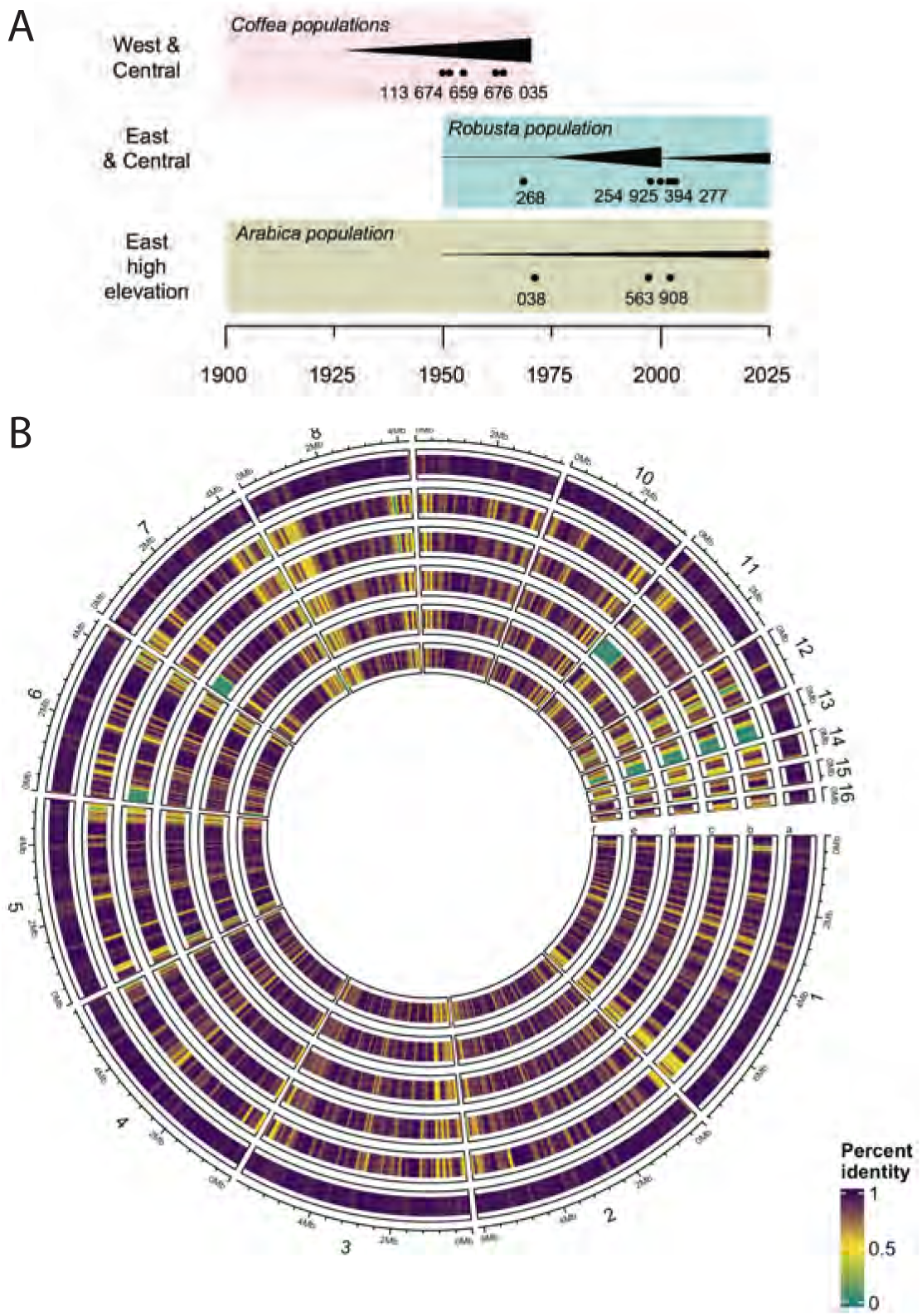
Diverse strain types within *Fusarium xylarioides*. **A** Schematic summary of geography and history of outbreaks and sampling times of isolates. Boxes = main coffee crops. Black triangles = qualitative incidence of coffee wilt disease. Dots = sample times for our isolates, labelled. Full strain details in Tables S1 and S2. Schematic not drawn to scale; incidence estimated from [10, 12, 14, 15]. **B** Selected *Fusarium xylarioides* genomes mapped to the *Fusarium xylarioides* arabica-specific reference genome. Comparison of contigs between the *Fusarium xylarioides* arabica563 reference genome and the selected *Fusarium xylarioides* genomes of the following host-specific strains: (a) Arabica038; (b) Robusta254; (c) Robusta268 (d) Coffea113; (e) Coffea035 (f) Coffea676. Regions with high coverage similarity to arabica563 are shown in purple; whilst absent regions in green. Each genomic window shown comprises 100kb with the consecutive window starting 25kb from the previous. The outer scale refers to contig size and the position of each genomic window.

**Figure 2:**
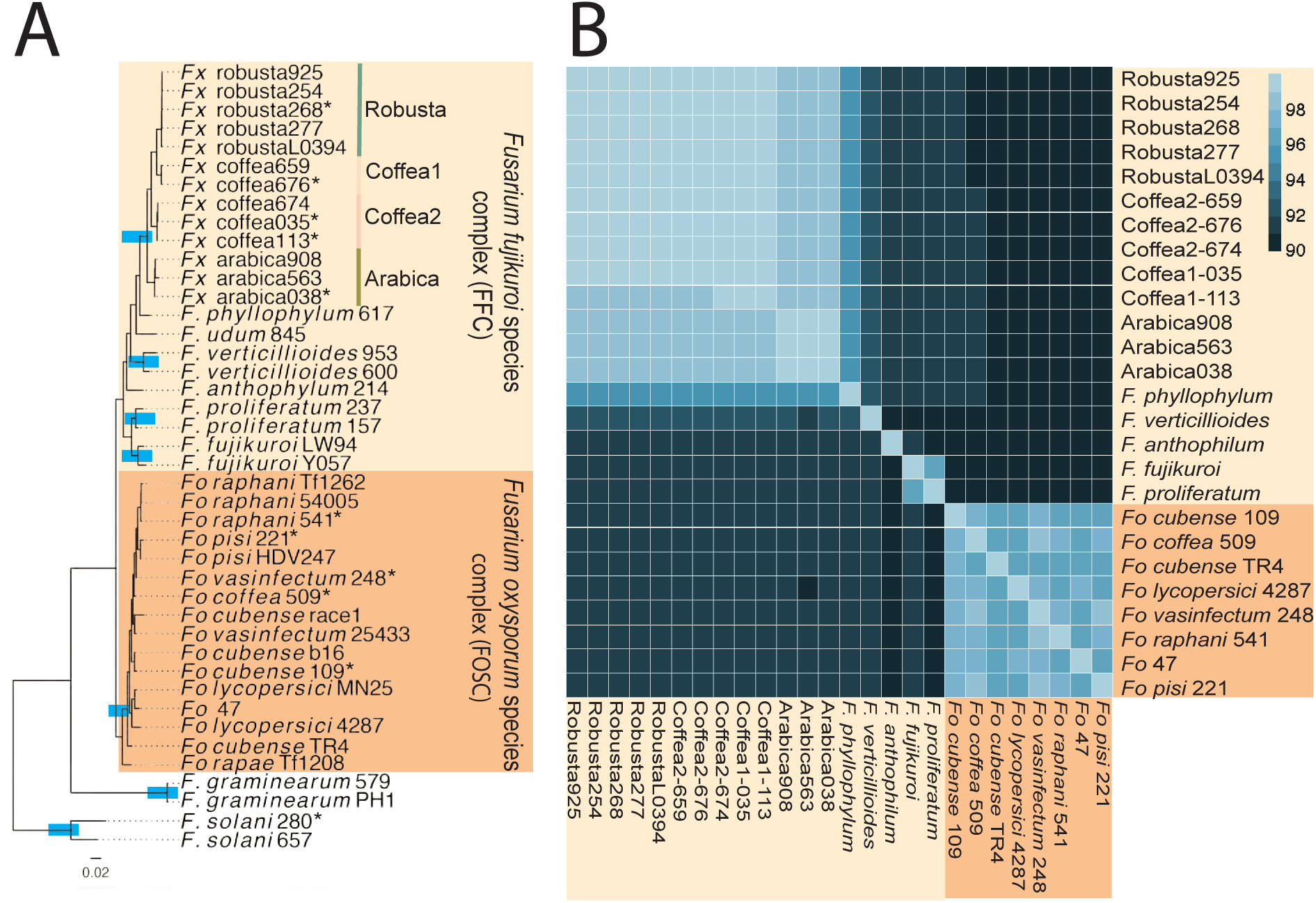
Phylogeny and whole-genome similarity between *Fusarium oxysporum* and *Fusarium fujikuroi* species complexes. **A** STAG species tree based on 3544 orthogroups in which every strain has at least one gene copy. Genomes sequenced in this study are indicated with an asterisk and blue boxes indicate resolved monophyletic species clades identified based on multilocus species delimitation. Shaded boxes refer to the the *Fusarium oxysporum* species complex (orange) and the *Fusarium fujikuroi* species complex (yellow), of which *Fusarium xylarioides* is a member. The *Fusarium xylarioides* populations arabica, robusta and coffea1 and coffea2 are labelled. Abbreviations: *Fusarium oxysporum, Fo*; *Fusarium xylarioides, Fx*. All strain details in Tables S1 and S2. **B** Whole-genome similarity. Cells indicate with colour the nucleotide similarity across all predicted coding sequences between the strains on the x and y axes. Shaded boxes refer to: the *Fusarium oxysporum* species complex, orange; and the *Fusarium fujikuroi* species complex, yellow, of which *Fusarium xylarioides* is a member. Arabica, robusta, coffea1 and coffea2 genomes all belong to *Fusarium xylarioides*. Excluding *F. xylarioides*, one genome is shown for each species/ *formae speciales* with strain details in Table S2. Abbreviations: Fo, *F. oxysporum*.

### The *Fusarium xylarioides* host-specific strains from different outbreaks belong to distinct populations

Analyses of genetic variation among *F. xylarioides* strains revealed population subdivision consistent with the hypothesis that multiple outbreaks arose from separate gene pools. Phylogenomics grouped the robusta strains into a monophyletic group, nested within the coffea strains from the first outbreak (1920s-1950s), and the arabica strains formed a distinct sister clade (Figure 3A). The coffea strains fall into two distinct clades, with coffea659 and coffea676 strains most closely related to the robusta group. Interestingly, coffea clade 1 contains coffea659, the only strain known to infect both arabica and robusta coffee trees [28] (host specificity for the other coffea strains has not been tested to our knowledge). A Neighbour-Net analysis based on 239,944 single nucleotide polymorphisms (SNPs) from coding and non-coding regions confirms the existence of four differentiated populations (Figure 3B): an arabica group, a robusta group and the same two coffea clades as recovered on the species tree (Figure 2A). While the robusta group shows a small amount of reticulation and therefore gene flow with the group containing coffea659 and coffea676 (Figure 3B), there is no reticulation either between or within the arabica and other coffea group, suggesting a lack of gene flow and recombination between and within the populations. Genetic differentiation between all four identified populations was further confirmed by high differentiation index values (F_ST_ all *>*0.96, absolute divergence d_xy_ all *>*0.0014, Table S3). Relationships between the four populations were not highly congruent across gene trees, e.g. *<*24% of single-locus gene trees supported the most common node between robusta and both coffea populations; and robusta and coffea-1 (arrows in Figure 3A). This lack of congruence may reflect incomplete lineage sorting due to short internode intervals. Internal nodes within each of the four populations had higher congruence, especially in the arabica and coffea clades (*>*40% of gene trees support the most common relationship). Such high levels of within-population genealogical congruence among loci could be due to high frequency of clonal reproduction.

**Figure 3:**
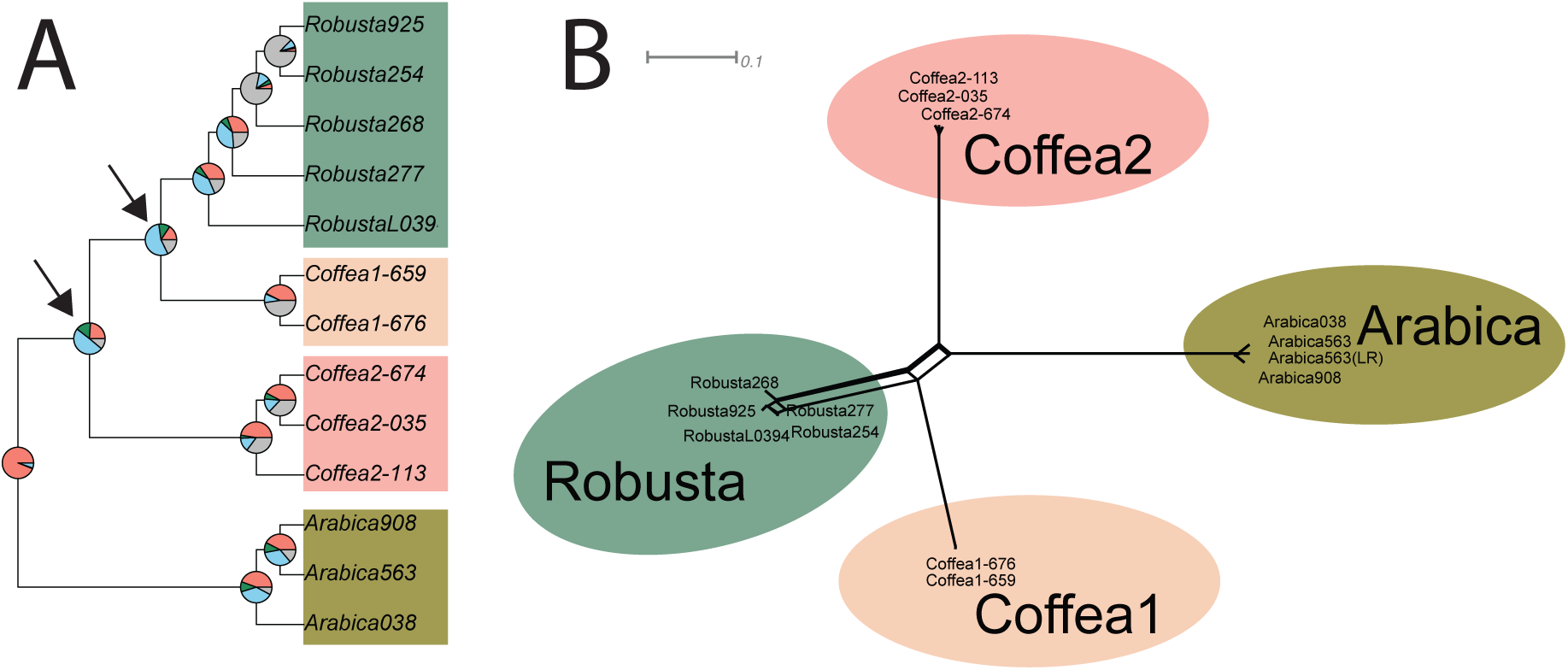
Genetic clusters in *Fusarium xylarioides*. **A** ASTRAL species tree in which phylogenetic relationships between *Fusarium xylarioides* genomes reconstructed from 1685 single copy orthogroups support monophyly of the clade, with little consistent support for alternate topologies. Full strain accession numbers are in Table 2. Pie chart colours: pink = proportion of genes(orthogroups) recovering the depicted node; dark green = the proportion of genes recovering the second most common topology; light blue = the proportion of genes recovering all other topologies; grey = unresolved topology. Arrows indicate the nodes connecting robusta with the coffea1 and coffea2 populations. **B** A neighbour-net (SplitsTree) analysis based on a single nucleotide polymorphism distance matrix. The scale bar represents 0.01 substitutions per site for branchlengths. The points are shaded according to their host population.

### Multiple large regions are shared between *Fusarium xylarioides* and different *Fusarium oxysporum* formae speciales

By comparing different *F. xylarioides* strains, we identified numerous large arabica-specific genomic regions were identified in the new arabica563 reference genome (Figure 1), representing putative HTRs that matched to effector genes and transposable elements identified from mobile and pathogenic *F. oxysporum* chromosomes, or to *Starships* (Figure 4). We identified an additional HTR in the robusta genomes that also contained matches to *F. oxysporum*’s pathogenic chromosome (Figure S2). HTRs present in both *F. xylarioides* and *F. oxysporum* but absent from more closely related species could potentially be involved in the wilt disease that both pathogens induce in their hosts.

**Figure 4:**
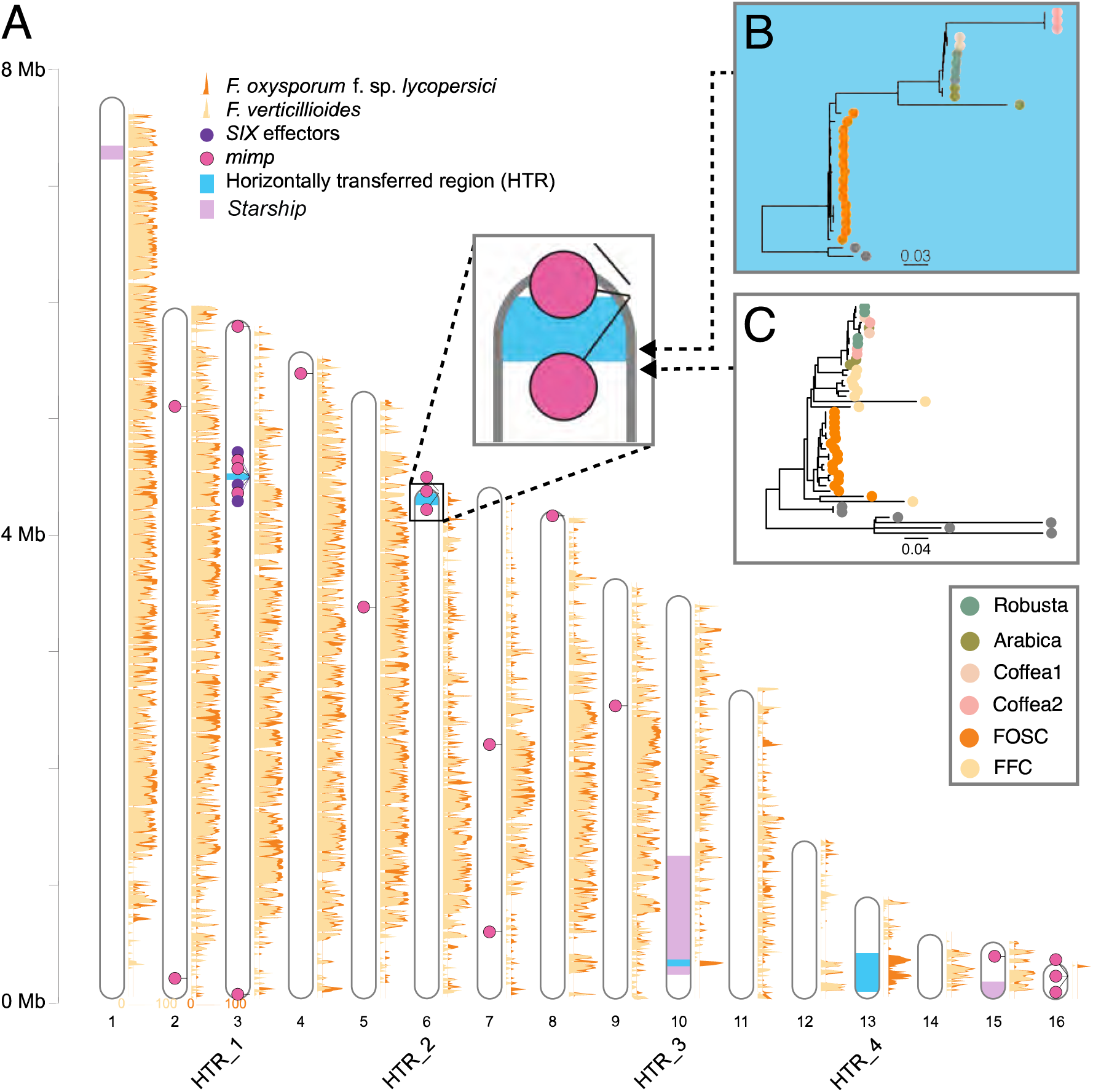
Similarity between the *Fusarium xylarioides, F. verticillioides* and *F. oxysporum* forma specialis *lycopersici* reference genomes and location of the horizontally transferred regions (HTRs). **A** The *F. verticillioides* and *F. oxysporum* f. sp. *lycopersici* long read assemblies were mapped to *F. xylarioides* in 10kb windows with no slide, with average similarity for each 10kb shown in yellow on orange polygons with *F. verticillioides* superimposed on *F. oxysporum* and orange shifted 0.2x right for visibility. Similarity for each is shown in legends on contigs 1 and 2. Only arabica563 contigs longer than 100 kb are shown. *SIX* effectors and *mimps* identified in the *F. xylarioides* reference genome are annotated, as well as each HTR which are labelled underneath the contig number. Genome accessions: *F. verticillioides*, accession AAIM02000000; and *F. oxysporum* f. sp. *lycopersici*, accession AAXH01000000. **B** A DendroBLAST rooted gene tree for the arabica563 gene P1J47 004978, found in HTR 2 region at 241 kb. Tip labels are colour coded by phylogenetic group, with FOSC representing the *F. oxysporum* species complex, and outgroup *F. solani* shown by grey tip labels. **C** A DendroBLAST rooted gene tree for the arabica 563 gene P1J47 004966, found in the HTR 2 flanking region at 259 kb. Tip labels are colour coded by phylogenetic group, with FOSC representing the *F. oxysporum* species complex and FFC the *F. fujikuroi* complex, and outgroup species *F. graminearum* and *F. solani* shown by grey tip labels.

All putative HTRs were absent from *F. verticillioides* as well as from other members of the *Fusarium fujikuroi* species complex (e.g. HTR 2 shown in Figure 4B-C). The HTRs range from small (5kb, HTR 1) to large (280kb, HTR 4), and all only return significant BLAST hits in public databases against *F. xylarioides* strains or various *F. oxysporum* formae speciales. HTRs were differentially present across the *F. xylarioides* populations (Figure 1): HTR 1 was found only in arabica and coffea populations, HTR 2 in all strains except robusta268 (the only pre-1970s outbreak robusta strain isolated), HTR 3 in both robusta and coffea populations but divergent between them; HTR 4 only in the arabica population, and HTR 5 only in robusta and coffea populations and most similar between robusta and coffea1 (Figure S4). We identified 133 coding genes in HTRs 1-4 in the arabica563 strain, and 10 coding genes in HTR 5 in the robusta strains. Of the arabica HTR genes, 103 had orthologs across all strains analysed in OrthoFinder. Of these orthologous groups, 36% were absent from the *F. fujikuroi* species complex, and among those which were present, the *F. xylarioides* arabica gene copies were closer to various *F. oxysporum* ff. spp (average phylogenetic distance was 0.13) than to those of various *F. fujikuroi* complex species (average distance was 0.33). In contrast, out of the 126 genes within the flanking regions 50kb either side of each HTR, nearly 90% of genes had orthologs that were similar between *F. xylarioides* arabica gene copies, and various *F. oxysporum* and *F. fujkuroi* complex species (0.24 and 0.22 respectively). The relationship observed among the flanking regions of the HTRs between *F. xylarioides* and the *F. oxysporum* and *F. fujikuroi* species complexes was the same as in the species tree (average phylogenetic distance was 0.09 and 0.07 respectively, Figure 2A), whereas the relationship observed inside the HTRs showed *F. xylarioides* and *F. oxysporum* more closely related.

HTR 1 comprises a 5kb region on contig 3. It contains three genes shared by all *F. xylar-ioides* arabica and coffea populations only (arabica563 gene copies are P1J47 016558, P1J47 017194, P1J47 017193) and which match to *F. oxysporum* effector proteins: Six7 with 91% identity; Six10 with 92% identity and Six12 with 94% identity (Table S5). Other *SIX* effector genes (*six1, six2, six6, six7* and *six11*) were identified in members of the *F. fujikuroi* species complex, with vertical (for *six2)* and horizontal inheritance from the *F. oxysporum* species complex suggested [29]. In addition, we previously identified *six7* and *six10* in arabica and coffea populations [26]. There are no significant BLAST hits in the intergenic regions of HTR 1 outside *F. xylarioides*. Each *SIX* gene either overlaps with a *mimp* (*six10* and *six12*, or includes a *mimp* in its promoter region (*six7*, *mimp* 943bp upstream). In *F. oxysporum* f. sp. *lycopersici*, these three genes are found on the supercontig 51 on the mobile, pathogenic chromosome, a region enriched in effector genes and *mimps*, among other transposable elements [30]. Remarkably, HTR 5, in the robusta and coffee populations, is shared with a different part of supercontig 51 to that shared with HTR 1 (Figure S2). In each genome, HTR 5 comprises a single scaffold (approximately 20 kb): All genes in HTR 5 are unique to the *F. xylarioides* robusta and coffea populations and various *F. oxysporum* ff. spp., as identified by BLAST. This region does not contain *SIX* effectors, but other types of genes important in infection, including a glycosyltransferase, Cytochrome P450 monooxygenases, a squalene-hopene-cyclase, a methyltransferase-UbiE protein and a Tri7 homolog.

The largest HTR (HTR 4) is on contig 13 in the *F. xylarioides* arabica strains. One half of contig 13 is found in all strains (e.g. Figure S3A), albeit with very low sequence similarity outside *F. xylarioides*. The other half is unique to the arabica population of *F. xylarioides* and *F. oxysporum* and shows low query coverage (*<*50% match to query length, BLAST) to the *F. oxysporum* genomes sequenced in this study, but high similarity to *F. oxysporum* strain Fo5176 (Accession CP128295.1, 94% similarity over 72% length). Compared to the best assembled *F. oxysporum* f. sp. *lycopersici* genome, a synteny alignment reveals that contig 13 matches to each of *F. oxysporum* f. sp. *lycopersici*’s mobile chromosomes, namely chromosomes 1, 3, 6, 14 and 15 (Figure S4). Within HTR 4, there are 39 genes with a high sequence identity between the *F. oxysporum* f. sp. *lycopersici* and *F. xylarioides* arabica strains (BLAST, *>*94% identity compared with 88% identity in flanking regions and whole genome identity of 90%, Figure S5D). Two of these genes belong to orthologous groups found just in *F. oxysporum* and the *F. xylarioides* arabica strains with highly similar *F. xylarioides* arabica copies and nested within diverse *F. oxysporum* copies from various *formae speciales* (Figure S5B-C).

Three *Starships* were identified in the arabica563 reference, and all were absent from at least one genome in those sequenced in this study and public genomes. Each *Starship* was bounded by flanking target site duplications with a predicted gene containing a DUF3435 domain (Pfam accession PF11917) located at the 5’ end of the element in the 5’–3’ direction (Figure S6). The *Starship* on contig 1 was in *F. xylarioides* and *F. phyllophylum* only, while those on contig 10 and 15 were only in the *F. xylarioides* arabica population, with the contig 10 *Starship* notably overlapping with HTR 3 (Figure 4A).

### Carbohydrate-active enzymes are highly expressed during infection and several have close matches in *Fusarium oxysporum*

Analysis of upregulated genes *in planta* during infection supported a role for putative HGTs in host adaptation, through the presence of effector genes. Diverse CAZyme families capable of breaking down cellulose and pectin from plant cell walls were previously identified in *F. xylarioides* arabica and robusta as putative effectors [26], but their role in infection remained unexplored. Analysis of gene expression of two strains of the *F. xylarioides* arabica clade from infected arabica coffee plants compared with the same strains grown *in axenic* liquid culture, revealed significant upregulation *in planta* of genes encoding CAZyme families important in the breakdown of pectin, xylan and other plant cell wall polymers (Figure 5A-B). In both strains, seven pectin lyases (within the polysaccharide lyase PL family) were predicted as effectors: three PL1 genes; three PL3 genes including the *pelA* and *pelD* effectors characterised by [31]; and one PL9 gene (Table S6). In both strains, most of the 23 pectin lyase genes from the subfamilies PL1, PL3, PL4, PL9, and PL26, showed strong up-regulation with 80% differentially expressed and 50% making up the top 20% of expressed genes (see Tables S9 and S10 for details). It appears that a subset of CAZyme families have been horizontally acquired in the *F. xylarioides* arabica population. *F. xylarioides* arabica563 was phylogenetically closer to various *F. oxysporum* formae speciales based on the genealogies of all 556 CAZyme orthologs (Figure 5C), compared to the phylogenetic distance in the species tree (Figure 2A). Other members of the *F. fujikuroi* species complex show similar phylogenetic distances for CAZyme and species tree genealogies (Figure 5C). Three CAZyme families were phylogenetically closer between *F. xylarioides* arabica563 and *F. oxysporum*: glycosyltransferases (GT, involved in the biosynthesis of polysaccharides), PLs and the associated carbohydrate binding molecules (CBM, which bind CAZymes to their substrate) (Figure 5D).

**Figure 5:**
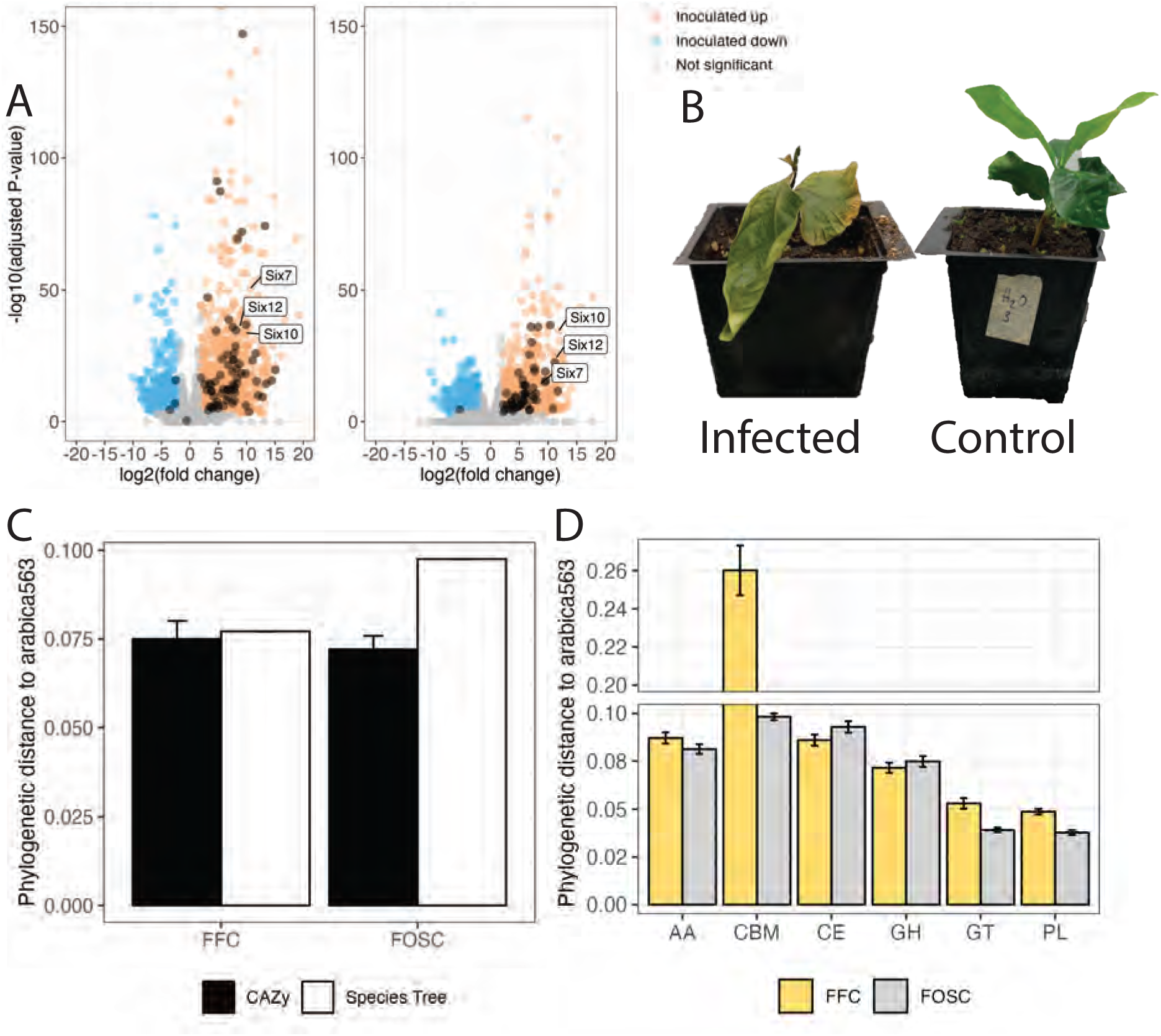
Up-regulated genes in *Fusarium xylarioides* coffee wilt infection appear to be horizontally transferred from *Fusarium oxysporum*. **A** Up-regulation of the *Fusarium oxysporum SIX*, that is secreted in xylem, effectors (annotated), and pectate lyase and other CAZymes involved in pectin metabolism (shaded dark orange). Significantly up-regulated *in planta* sample genes were identified using negative binomial generalized linear models and shaded orange and those down-regulated are shaded blue. The CAZymes encompass AA9, CE12, CE5, GH105, GH11, GH12, GH43, GH5, GH51, PL1, PL3, PL4, PL9, PL26. **B** Arabica coffee plants 99 days post inoculation, infected at left, that is inoculated with *Fusarium xylarioides* arabica563, and control at right, that is mock-inoculated with sterile water. **C-D** Certain *Fusarium xylarioides* CAZyme families are more closely related to *Fusarium oxysporum* than to the *Fusarium fujikuroi* species complex. **C** Comparing phylogenetic distances between *F. xylarioides* arabica563 with the *F. fujikuroi* species complex (FFC); and *F. xylarioides* arabica563 with the *Fusarium oxysporum* species complex (FOSC) for CAZymes and species tree gene families, reveal that CAZymes are more closely related for the *Fusarium oxysporum* species complex, whilst there is no difference for the *F. fujikuroi* species complex. CAZyme gene family number shared with *F. xylarioides* arabica563 = 525 for the *F. fujikuroi* species complex; 543 for the *F. oxysporum* species complex. Error bars show standard error. **D** Gene families that match CAZYme glycosyltransferases (GT), pectin lyases (PL) and the associated carbohydrate binding modules (CBM), are phylogenetically closer between *F. xylarioides* arabica563 and the *F. oxysporum* species complex (grey bars, FOSC) than with members of the *F. fujikuroi* species complex (yellow bars, FFC). There is no difference for CAZyme auxiliary activity (AA), carbohydrate esterase (CE) nor glycoside hydrolase (GH) gene families. Error bars show standard error.

### Effectors known from *Fusarium oxysporum* mobile chromosomes are highly expressed during coffee wilt infection

Several previously identified putative effectors [26] were also found to be significantly upregulated in *F. xylarioides* from infected coffee plants compared to fungi grown in culture (Tables S7 and S8). Six putative effectors known from the mobile pathogenic chromosome of *F. oxysporum* f. sp. *lycopersici* were up-regulated in at least one arabica strain. These were: P1J47 011695 named by its orthologous group and inferred to be a divergent type of *six7* effector (referred to as OG0014398 in [26]); three novel effector candidates identified by [30] which are all secreted proteins - unidentified *FOXG 14254*, oxidoreductase *orx1* and a catalase-peroxidase found in infected xylem sap *FOXG 17460*; *sge1*, the *SIX* effector gene transcription factor; P1J47 017133, a LysM effector (referred to as OG0013477 in [26]; and two pectin lyase effectors, *pelA* and *pelD*. Only one putative effector, P1J47 017133, was expressed less in infected plants than control *in axenic* culture samples across both strains. Seven remaining putative effectors were expressed but not differentially in both strains and six genes were expressed but not differentially in one strain (Tables S7 and S8). In addition to these putative effectors, reanalysis of the new reference genome with EffectorP 3.0 detected further pectinolytic enzymes, similar extracellular degradative enzymes, as well as the established fungal effector domains Ecp2 and LysM (see Table S6 for details), which were also upregulated in infected plants.

Of the 496 genes that were significantly upregulated *in planta* in both arabica strains, 19 were absent from *F. fujikuroi* complex species and mostly did not match any recognised domains (Table S5). Of these 19 genes, remarkably, three were found in HTR 1 and were shared by *F. xylarioides* arabica and coffea populations only (P1J47 016558, P1J47 017194, P1J47 017193). Each matches to an *F. oxysporum* effector protein: Six7, Six10 and Six12 respectively (Table S5). The *SIX* orthologues were also in the top 20% most highly expressed genes *in planta*, across the arabica strains (Figure 5B). Of the remaining genes absent from *F. fujikuroi* complex species, several were population-specific; some were shared by all *F. xylarioides* populations; and some were shared between *F. xylarioides* and other species complexes (Table S5).

### Miniature *impala* elements and other transposons are shared between *Fusarium xylarioides* and *Fusarium oxysporum*

The *F. xylarioides* genomes contained miniature *impala* elements (*mimps*) that are present in the more distant *F. oxysporum* genomes but lacking in more closely related species, as well as other transposons previously identified in mobile *F. oxysporum* chromosomes. These elements are present in some putative HTRs and close or overlapping to several highly expressed genes. A total of 450 full-length *mimps* were detected across the *F. xylarioides* and *F. oxysporum* genomes (Figure 6A). *Fusarium oxysporum* formae speciales genomes contained the most *mimps*, with *F. oxysporum* f. sp. *lycopersici* and *F. oxysporum* f. sp. *raphani* in the highest copy numbers (90 and 85 respectively, Figure 6). The *mimp* families 1, 2 and 4 were the most numerous in all genomes. Within *F. xylarioides*, robusta and coffea strains contained twice as many *mimps* as the arabica strains, primarily driven by *mimp* families 1 and 2. All *F. xylarioides* genomes and most of the *F. oxysporum* genomes contained intact *impala* transposases (Figure 6A), required for *mimp* activity [32]. No *mimps* or *impala* transposases were found in the genomes of other species in the *F. fujikuroi* species complex. Phylogenetic reconstruction showed that *mimps* in *F. xylarioides* are intermingled with those from the *F. oxysporum* species complex, rather than forming separate clades, consistent with either a frequent and ongoing transfer between the two species, or a single transfer of multiple diverse copies (shown for *mimp* family 1, Figure 6B).

**Figure 6:**
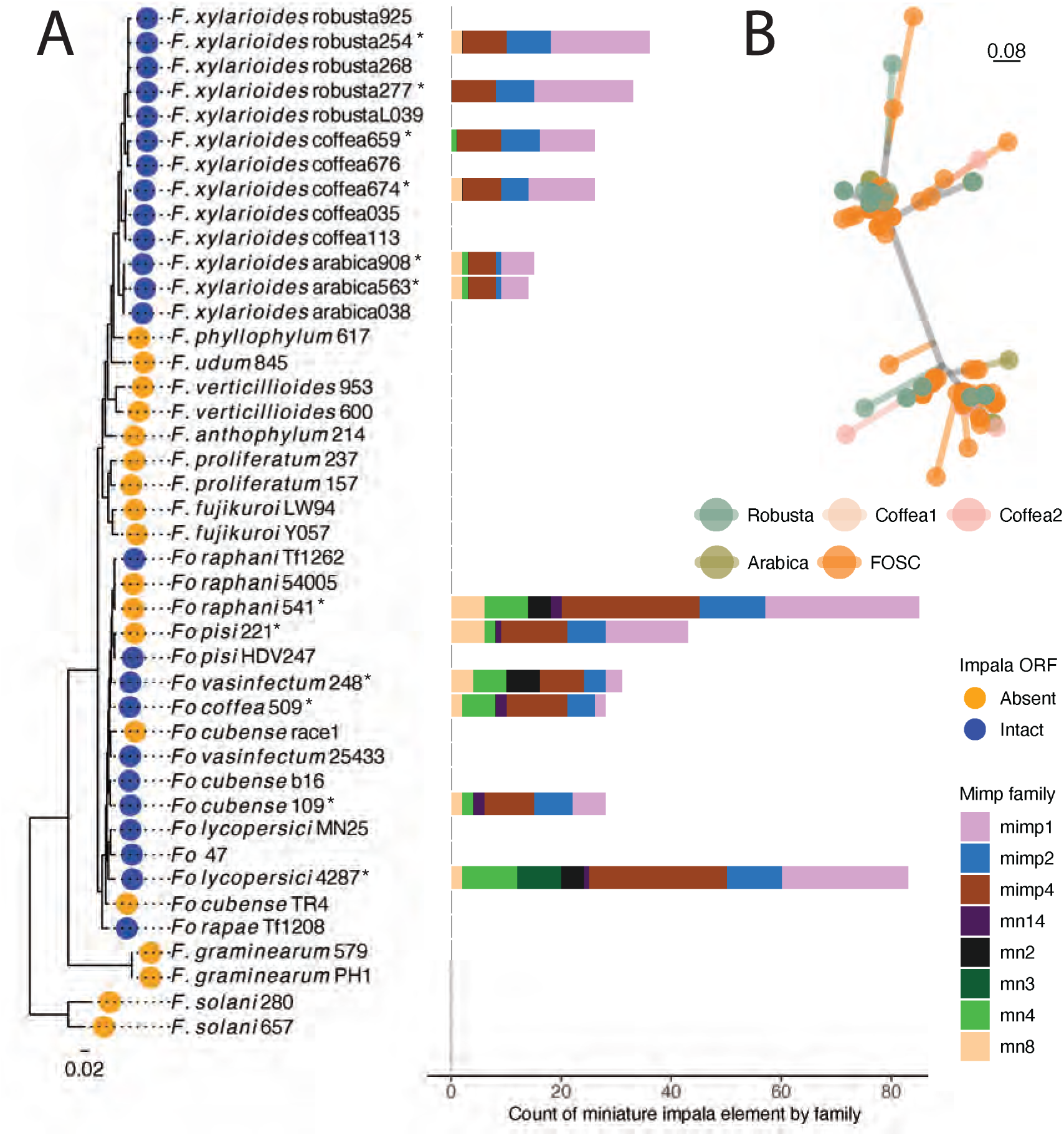
Miniature *impala* composition of *Fusarium* genomes. **A** *Miniature impala (mimp)* elements across the whole-genome are shown as a count per family. They were identified in six representative *Fusarium xylarioides* genomes, all *Fusarium oxysporum* formae speciales genomes sequenced in this study and *F. oxysporum* f. sp. *lycopersici*, each annotated strain denoted by an asterisk. Colour represents the family to which the *mimp* sequence belongs. The STAG species tree is shown at left and the tip label corresponds to the presence of an intact *impala* transposase open reading frame. **B** *Fusarium oxysporum* and *Fusarium xylarioides miniature impala (mimp)* family 1 sequence genealogy. All *mimp* 1 sequences have been extracted and aligned with macse [88], with phylogeny represented by an unrooted tree. Each point corresponds to a *mimp* sequence and the treescale shows their evolutionary distance. Tree tip labels are shaded by phylogenetic group: arabica, robusta, coffea1 and coffea2 *F. xylarioides* populations; FOSC, *Fusarium oxysporum* species complex. Full strain details in Table 2.

Some *mimps* were observed in large regions identified as putative HTRs above. For example, in regions that match *F. oxysporum* f. sp. *lycopersici*’s supercontig 51, *F. xylarioides* arabica contained at least 3 and robusta at least 2 *mimps*. Some *mimps* overlapped with (*six10* and *six12*) or were close to the *six7* effector genes (Figure S2A). The coffea scaffolds that matched the region of supercontig 51 found in *F. xylarioides* arabica also contained *mimps*, whilst those matching the region found in *F. xylarioides* robusta lacked these, and both robusta and coffea scaffolds contained the unnamed TE family rnd-6 family-1942 and newly-recognised TEs (Fot6, MGR583-like, Yaret1) found on *F. oxysporum* f. sp. *lycopersici*’s mobile pathogenic chromosome (identified by [30]). It is possible, however, that the *mimps* were assembled to different coffea scaffolds, as the robusta scaffolds that match supercontig 51 are only 20kb each.

In addition to *mimps*, HTRs 1-3 and 5 all contained diverse transposable elements which were first identified on the mobile pathogenic chromosome of *F. oxysporum* f. sp. *lycopersici* by [30] (Figure S7). Across the *F. xylarioides* arabica563 genome, TE density (excluding simple and low complexity repeats) was 0.009, whilst most HTRs contained more transposons (Table S12. The average across HTRs and *Starships* were 0.1 and 0.064 respectively. HTRs 1, 2 and 5 had the highest TE densities at 0.2, 0.09 and 0.09, with numerous (50%, 12%, and 83% respectively) matching to transposons originally identified on *F. oxysporum* f. sp. *lycopersici*’s mobile pathogenic chromosome [30] and the unnamed TE family rnd-6 family-1942 which we found with *mimps* (Figure S7).

## Discussion

We have discovered genetic subdivision in *F. xylarioides*, namely multiple differentiated host-specific populations isolated from different species of coffee. We found that these populations share large mobile *Starship* elements, as well as several highly similar regions with *F. oxysporum* that are lacking in other closely related species, as well as the, consistent with their acquisition by horizontal transfer. Transcriptome data following infection revealed high expression of previously identified effector genes, including some only known to *F. oxysporum* outside of coffee wilt, as well as pectinolytic enzymes. Finally, we found that all *F. xylarioides* populations share highly similar *mimps* and other transposable elements with *F. oxysporum* that have likely transferred between the species and could have facilitated HGT of wider genome regions.

The *F. xylarioides* arabica and robusta groups were genetically differentiated from each other and from coffea1 and coffea2, which were themselves two differentiated populations. The two coffea populations overlapped spatially and temporally (coffea 1 was isolated from Guinea and Central African Republic, coffea2 was isolated from Guinea, the Central African Republic, and Cote d’Ivoire), yet they remained genetically distinct, with the robusta population apparently emerging from within coffea 1. Possible causes of genetic differentiation could be the occurrence of distinct clonal lineages, consistent with the congruence between gene genealogies within populations, or reproductive isolation due to pleiotropy between host adaptation and mate choice [33]. The specific coffee tree hosts from which *F. xylarioides* coffea population strains were isolated were not recorded, so we do not know if these infected broad or specific coffee tree species. Reproductive isolation can be a pleiotropic effect of specialisation if mating occurs within the host [33], as only strains able to infect the same trees can mate. Indeed, mating appears to occur in the host in *F. xylarioides* as perithecia are only observed on coffee wilt diseased trees [34], which could have aided genetic differentiation between geographically overlapping populations. Our findings support earlier work based on DNA markers and crossing experiments [35, 36], which suggested that *F. xylarioides* was a species complex containing distinct arabica and robusta groups. The authors called the isolates *F. xylarioides* f. sp. *abyssiniae* and *F. xylarioides* f. sp. *canephorae*, respectively. Here, we confirm that: the *F. xylarioides* formae speciales proposed nearly two decades ago can now be recognised as genetically differentiated populations with different host-specificity.

Several lines of evidence favour the acquisition of large genome regions by horizontal transfer (HTRs) into *F. xylarioides* populations over alternative hypotheses. First, the alternative scenario of presence in the common ancestor and vertical inheritance is contradicted by a higher sequence identity with *F. oxysporum* than expected from other genes. Second, the population-specific nature of these regions, present in either the arabica or robusta populations and occupying mostly regions of the genomes which are absent from the other, and at different loci in genomes, rules out introgression by inter-specific sexual hybridisation. In contrast, the locations of HTRs are homologous between strains within each population, suggesting a single transfer event into a recent common ancestor for each population. Third, horizontal transfers can occur through transposon-assisted mechanisms or by transfer of an entire chromosome [21] and *Fusarium* species are known to form anastomoses - somatic fusions between mycelia. Such fusions may facilitate the transfer of genetic material by whole chromosomes [37]. Indeed, hyphal fusions can occur between the arabica and robusta *F. xylarioides* populations [38]. Finally, there is a lower sequence identity along these regions between *F. oxysporum* formae speciales genes than between gene copies found in *F. xylarioides*, in which each region is identical in each strain. However, the incomplete identity (*<*100%) with *F. oxysporum* suggests that the *F. oxysporum* f. sp. *lycopersici* genome is not the direct donor strain, the donor is from a distinct population, or the HTRs are more ancient than the outbreaks. The occurrence of horizontal transfer of pathogenicity genes via accessory chromosomes was first reported in *F. oxysporum* f. sp. *lycopersici* with horizontal transfer of its entire mobile pathogenic chromosome and also a large duplicated region from chromosome 3 to two previously non-pathogenic strains [21]. Several more cases have been reported since then, and each with a correlation with *mimps* in transposition [29, 30]. In the HTRs discovered in this study, a whole mobile chromosome may have been acquired by a common ancestor to the arabica and robusta populations, with later divergence and differential loss of genomic regions and integration of different parts. Alternatively, only parts of the *F. oxysporum* chromosome may have been transferred into *F. xylarioides*, with different parts in arabica and robusta populations. Gene loss of regions that are no longer useful to the organism is well reported [39].

The association of *mimps*, between *F. oxysporum* and *F. xylarioides*, sheds further light on potential mechanisms of horizontal transfer. Overall, we found four putative HTRs with transposable elements specifically linked to horizontal transfer in *F. oxysporum*. This suggests a role in HGT and generating effector gene diversity. This could comprise either direct transposition (as in HTRs 1-3 and 5) or via increasing the potential for recombination events (including ectopic recombination and genome loss). One HTR (4) lacked *mimps* and other transposons and could suggest a different transfer mechanism. We found no *mimps* in the *F. fujikuroi* complex species (although [29] reported a few copies in some species). The arabica and robusta *F. xylarioides* populations contained different numbers of *mimps* in total as well as in their HTRs; this is consistent with a different history of transfer. Such horizontal transfer events may have allowed the acquisition of several CAZyme gene families, important in vascular wilts; this may explain the greater similarity between *F. xylarioides* and *F. oxysporum* than other members of the *F. fujikuroi* species complex. In *F. oxysporum*, *six10, six12* and *six7* make up supercontig 51 and are flanked by *mimps* [30]. In the *F. xylarioides* arabica population, *six12* and *six10* contained *mimps*, while *six7* was *<*1 kb from one. A different piece of supercontig 51 is present in the robusta genomes, which also contained *mimps* and other class II transposable elements, as well as genes involved in pathogenicity. No other *SIX* mini-clusters were present or expressed in *F. xylarioides*. A study of two *F. oxysporum* f. sp. *cubense* transcriptomes found that both lack mobile chromosomes of *F. oxysporum* f. sp. *lycopersici*, yet still show differing presence and expression of *SIX* genes [40]. In *F. oxysporum*, all effector genes contain a *mimp* in the promoter region [30], yet in *F. xylarioides* this was only the case for the three *SIX* genes. In addition, the fact that the *mimp*-overlapping *SIX* genes are only found in the arabica and coffea populations (and not robusta, nor any *F. fujikuroi* complex species) suggests a horizontal transfer origin.

To our knowledge, such interspecies horizontal transfer of pathogenicity has not been reported in *Fusarium* before, while introgression has [41]. The close phylogenetic relationship between CAZyme gene familes from *F. oxysporum* and the arabica strains provides further evidence of horizontal transfer of gene types that are important in vascular wilt infection. Such horizontal transfer could have occurred in a shared niche, since both *F. xylarioides* and *F. oxysporum* are soil-borne pathogens and have been isolated from the roots and wood of coffee wilt-diseased trees in Ethiopia and central Africa [42], as well as from banana roots in an intercropped field in Uganda [43], and on which *F. oxysporum* f. sp. *cubense* is widespread [44]. These transfers may have driven the evolution of new host-specific populations and therefore could have facilitated disease emergence and subsequent re-emergence. Recently, studies have shown the value of comparative genomics in elucidating the role of these changes in domesticated fungi and plant pathogens, including a host jump in *Phytophthora infestans* followed by adaptive specialisation [45], transfer of a 14kb fragment encoding the ToxA virulence protein and abundant transposons between three wheat fungal pathogens [18], and the transfer of regions that allow adaptation to cheese environments in distantly related species [46]. Furthermore, transfer of pathogenicity into a previously nonpathogenic strain in *F. oxysporum* has been demonstrated experimentally [21]. These examples, as well as cross-kingdom transfers (e.g. [47], demonstrate the occurrence of HTRs across many distantly related branches of the eukaryotic tree of life and their importance for shaping eukaryotic evolution [48]. Previously, the analysis of historic genetic material involved the use of highly degraded museum specimens and molecular markers and target probes in a small number of genes [49]. However, fungal culture collections contain tens of thousands of fungal isolates collected over the past century and preserved in a living state [50]. If collections are connected with new -omics studies, new HTRs could be investigated at a new inter- and intra-species level.

## Materials and Methods

### Long-read reference genome sequencing and assembly

The *F. xylarioides* strain IMI 389563 (hereafter called “arabica563”) from the arabica host-specific strain group of coffee wilt disease was used for long-read sequencing, and is available from the CABI-IMI culture collection (CABI, Egham, UK). Arabica563 mycelium was grown in GYM broth for 7 days at 25°C and 175 RPM. High-quality high molecular weight genomic DNA was extracted from 1-2g of mycelium using the chloroform extraction method in [51] (full details in supplementary methods). The gDNA quantity was determined using a NanoDrop 2000 (ThermoFisher) and its integrity was determined by electrophoresis with a 0.5% agarose gel. 1 *µ*g of gDNA was diluted with nuclease-free water to a final volume of 47 *µ*l. Half of the DNA was then sheared into 20 kb fragments with a COVARIS g-TUBE (Covaris, Inc., Woburn, MA, USA), centrifuging at 4,200 rpm for 1 minute to increase DNA throughput. The shared DNA was converted into Oxford Nanopore libraries using the SQK-LSK110 library kit and the short fragment buffer (SFB) (Oxford Nanopore Technologies, UK). Library prep followed the standard manufacturer’s protocol apart from the final elution, which was done for 30 minutes at 37*^o^*C. The libraries were sequenced using a SpotON R9.4.1 FLO-MIN106 flowcell for 48 hours.

Nucleotide bases were called from the raw sequence data using the super high accuracy model (SUP) of the GPU version of Guppy v6.0.1 (config file: dna r9.4.1 450bps sup.cfg). Base-called reads were assembled into contigs by Flye v2.9 with “-nano-hq” parameters [52]. The Flye assembly was error-corrected using both Nanopore and Illumina data with one round of medaka v1.4.4 (https://github.com/nanoporetech/medaka) and one round of (Hapo-G) polishing [53]. Contig completeness was visualised in Tapestry v1.0.0 [54], including the presence of the terminal telomere repeat sequence TTAGGG, common to eukaryotes.

### Illumina sequencing and assembly

An additional 11 *Fusarium* genomes were selected for Illumina whole-genome sequencing: five *F. xylarioides*; five *F. oxysporum* and one *F. solani* (Table 2), see supplementary methods for strain details. Of the five *F. xylarioides* strains, three were from the pre-1970s outbreak, which was observed to infect multiple *Coffea* species and which we call “coffea” (coffea035, coffea113, coffea676), and the remaining two were the earliest isolates available in the collections for each of the arabica and robusta groups (both isolated from around 1970, arabica038 and robusta268).

Strain morphologies were verified and grown in GYM broth following the same protocol as above. DNA was extracted from 250-500mg of washed mycelium following the NucleoSpin Soil Mini Kit (Macherey Nagel, Germany) standard protocol. For each strain, a single library was prepared with the Illumina DNA PCR-Free Prep and sequenced with the Illumina Novaseq 6000 with 2x 150bp reads. For details of assembly and quality checking, see supplementary methods.

Each *F. xylarioides* MEGAHIT assembly, see supplementary methods for strain details, was mapped to the reference genome assembly using RagTag v2.1.0 homology-based scaffolding [55]. The parameter “-u” identified unmapped (and therefore absent from arabica563) scaffolds. The *F. oxysporum* and *F. solani* MEGAHIT assemblies were mapped to their closest and least fragmented publicly available assembly (Table 1). The *F. oxysporum* f. sp. *lycopersici* and *F. oxysporum* f. sp. *cubense* genomes were the least fragmented (see N50 values). These RagTag assemblies were used to predict transposable elements (see later section).

**Table 1:** Genomes sequenced in this study. The columns describe: species, type of sequencing completed, RNA-seq data used as evidence in gene prediction, expressed sequence tag (EST) data used as evidence in gene prediction, whether the genome was used in Nsimscan for whole-genome similarity, which publicly available assembly the genome was scaffolded to using RagTag.

| Species | IMI Strain | Funannotate:<br>RNA_Seq data | Funannotate:<br>EST data | RagTag scaffolded to |
| --- | --- | --- | --- | --- |
| *F. oxysporum*<br>f. sp. *cubense* | 141109 | ERR10015931 | *F. oxysporum* f.sp.<br>*cubense* II5 v2.0<br>[59] | GCA 005930515.1 |
| *F. oxysporum*<br>f. sp. *coffea* | 244509 | SRR10123445 | *F. oxysporum* f.sp.<br>*cubense* II5 v2.0<br>[59] | GCF 000149955.1 |
| *F. oxysporum*<br>f. sp. *vasinfectum* | 292248 | SRR12855423 | *F. oxysporum* f.sp.<br>*cubense* II5 v2.0<br>[59] | GCA 000260175.2 |
| *F. oxysporum*<br>f. sp. *raphani* | 337541 | SRR14318405 | *F. oxysporum* f.sp.<br>*cubense* II5 v2.0<br>[59] | GCA 019157275.1 |
| *F. oxysporum*<br>f. sp. *pisi* | 500221 | SRR12855423 | *F. oxysporum* f. sp.<br>*pisi* F23 v1.0 | GCA 000260075.2 |
| *F. solani* | 392280 | SRR19182467 | *F. solani* FFSC 5 v1.0<br>[60] | GCA 020744495.1 |

The raw sequence data and assembled genomes have been deposited in the International Nucleotide Sequence Database Collaboration (INSDC) database under the BioProject accession number PRJNA1043203.

### RNA sequencing to aid annotation of long- and short-read assemblies

The same arabica563 strain used for long-read sequencing was grown under the same culture conditions described above. Mycelia were harvested and total RNA extracted using the standard protocol of the RNeasy Mini kit (Qiagen, Hilden, Germany). The RNA quality assessment (Qubit and Fragment Analyzer), library preparation and Next Generation Sequencing (60M paired-end reads per sample) were performed by Genewiz/ Azenta (Frankfurt, Germany), see supplementary methods for QC steps. Transcript sequences were *de novo* assembled using Trinity v2.13.2 with the genome-free method [56].

### Gene prediction and annotation

Protein-encoding genes were predicted in all short-read genomes using the Funannotate Eukaryotic Genome Annotation Pipeline v1.8.11 [57]. All 13 *F. xylarioides* genomes were annotated (Table 2), including re-annotations for the six genomes from [26]. Before annotation, repetitive contigs were removed using “funannotate clean” with default parameters and repetitive elements within the assemblies were soft masked using RepeatModeler and RepeatMasker with “funannotate mask” and *F. oxysporum* parameters.

**Table 2:** Statistics for sequenced *Fusarium* genomes. Arabica563 is the reference assembly. Genes were not predicted on the arabica563 reference assembly. Only contigs *>*1kb in length were included. *Fusarium oxysporum* is abbreviated to Fo.

| Short name | Strain ID (IMI) | Genome size Mb (>1kb) | No. contigs (>1kb) | N50 kb | Complete BUSCO % | GC % | Coding Genes |
| --- | --- | --- | --- | --- | --- | --- | --- |
| Arabica563 | 389563 | 57.3 | 21 | 5,156 | 100.0 | 42.2 |  |
| Arabica038 | 507038 | 51.1 | 4631 | 24.5 | 97.9 | 43.4 | 15,705 |
| Robusta268 | 392268 | 50.2 | 4420 | 27.1 | 97.6 | 44.1 | 15,575 |
| Coffea035 | 507035 | 51.1 | 3880 | 30.4 | 97.9 | 43.9 | 15,588 |
| Coffea113 | 507113 | 48.7 | 3795 | 30.1 | 98.3 | 44.3 | 15,115 |
| Coffea676 | 392676 | 50.0 | 4362 | 27.1 | 97.9 | 44.4 | 15,649 |
| *F. solani* 280 | 392280 | 48.4 | 1867 | 45.2 | 96.9 | 50.9 | 14,007 |
| *Fo coffea* 509 | 244509 | 49.7 | 1923 | 67.9 | 98.3 | 47.4 | 14,475 |
| *Fo cubense* 109 | 141109 | 48.4 | 1992 | 60.1 | 99.0 | 47.6 | 14,993 |
| *Fo pisi* 221 | 500221 | 52.9 | 4299 | 28.1 | 95.5 | 47.5 | 16,614 |
| *Fo raphani* 541 | 337541 | 53.8 | 4285 | 38.1 | 98.3 | 47.7 | 16,421 |
| *Fo vasinfectum* 248 | 292248 | 49.4 | 1904 | 72.5 | 98.6 | 47.4 | 14,846 |

The trimmed RNA-seq reads and the trinity assembly were aligned to the arabica563 MEGAHIT assembly using the parameters “funannotate train –jaccard clip” which runs PASA to model the gene structures. The PASA gene models were parsed to run Augustus, snap, GeneMark and GlimmerHMM, with the predictions used to run CodingQuarry and the Funannotate Evidence Modeler. The complete set of arabica563 output files were then used to predict protein-encoding genes on the remaining MEGAHIT genome assemblies using “funannotate predict -p predict results/fusarium xylarioides 389563.parameters.json”. The arabica563 MEGAHIT assembly was selected over the reference genome to avoid issues with gene prediction on assemblies of different quality when the parameters were applied to the remaining *F. xylarioides* MEGAHIT assemblies. The gene model predictions were refined (e.g. correcting intron-exon boundaries) and untranscribed regions annotated by re-aligning the RNA-seq data for each genome using “funannotate update” on the output files from “funannotate predict”.

The annotations were then mapped to the long-read RagTag assemblies using Liftoff [58] with default parameters.

### Other *Fusarium* genomes

The same steps were followed for the five *F. oxysporum* genomes and the *F. solani* genome, using different sources of evidence for the different species. Sorted BAM files (RNA-Seq reads for the closest *f. sp.* match were downloaded from the sequence read archive and aligned to the corresponding assembly using “bbmap.sh”), and expressed sequence tag data for the closest *f. sp.* match from JGI Mycocosm were used to provide evidence to “funannotate predict”. The expressed sequence tag data were produced by the US Department of Energy Joint Genome Institute https://www.jgi.doe.gov/ in collaboration with the user community. Evidence sources are listed in Table 1. The same RNA-Seq data was then re-aligned through “funannotate update” as in the section above.

Functional annotation of the protein-coding genes for all genomes was completed with annotations from PFAM, InterPro, UniProtKB, MEROPS, CAZyme and GO Ontology through “funannotate annotate”. InterProScan v5.56-89.0 was run locally using the parameters “-goterms -iprlookup” and phobius was run through the web browser (https://phobius.sbc.su.se/). All proteins lacking significant hits were annotated as hypothetical proteins.

### Transposable elements and repeat annotations

Repeats and transposable elements (TEs) were identified from the nucleotide assemblies. A custom repeat library was constructed using all *Fusarium* RagTag assemblies in RepeatModeler v2.0.3 with the parameter “-LTRStruct” [61]. Added to this library were TEs known to be involved in *F. oxysporum* transfer of pathogenicity between strains: *impalas*, a new set of TEs described by [30], as well as manually curated *mimps*. Sequences for *mimp* families 1-4 were downloaded from NCBI (accession numbers: AF076624.1; AF076625.1; EU833100.1; EU833101.1). Sequences for the more divergent *mimp* families mimp5, mimp6, mn2, mn3, mn4, mn8 and mn14 are from [32]. Manual curation followed the protocol described by [62] and involved: identifying conserved domains with sequence homology to these TEs using BLAST; only retaining the best BLAST hits (*>*= 50% query length and *>*= 80% identity) for each TE family; extracting and aligning nucleotide sequences for TE families including an additional 500 bp up- and down-stream; trimming alignments to the TIRs at their start and end and removing insertions using T-COFFEE [63]. This generated consensus sequences for each *mimp* family that were added to the RepeatModeler custom library and used to call TEs and repeats from genome scaffolds using RepeatMasker v4.1.2 with the parameters “-lib $LIB -gff -no is” [64]. The presence of the *impala* transposase ORF from *F. oxysporum* f sp *melonis* was found by a BLAST search “tblastn -evalue 1.00e-5” using accession AAB33090.2.

### Orthologous gene groups, species tree and genetic structure

Whole-genome similarity between the protein-encoding genes was assessed using NSimScan, part of the QSimScan package [65]. OrthoFinder v2.5.4 was used to determine *Fusarium* orthologous groups between the genomes sequenced in this study (Table 2) and published genomes detailed in Table S2 [66], see supplementary methods for full details. The species tree was built and rooted using STAG in OrthoFinder [67, 68] with 3544 orthogroups in which every strain has at least one gene copy. Resolved monophyletic species clades were identified using tr2 software [69]. Concordance of the set of individual gene trees for single ortholog group nucleotide sequences was evaluated using a species tree built in ASTRAL [70] and summarised using PhyParts (https://bitbucket.org/blackrim/phyparts/src/master/).

Single nucleotide polymorphisms (SNPs) were called against the arabica563 reference genome for each set of *F. xylarioides* reads, see supplementary methods for strain details. SNPs were called using GATK v4.1.2.0 “HaplotypeCaller -ERC GVCF”, which provides one genomic variant call format (gVCF) output file per strain. Using Snakemake v5.3.0, gVCFs were combined using GATK “CombineGVCFs”, genotypes were combined with “GATKGenotypeGVCFs”, “SelectVariants -select-type SNP” selected the SNPs which were then filtered with “VariantFiltration QUAL*<*30, DP*<*10, QD*<*2.0, FS*>*60.0, MQ*<*40.0, SOR*>*3.0, QRankSum*<*-12.5, ReadPosRankSum*<*-8.0”. The SNP dataset was used to infer population structure. Principal component analysis of SNP data was completed using the R packages SNPArray and gdsfmt with a linkage disequilibrium threshold of 0.05 [71, 72]. The nucleotide diversity *π*, Watterson’s *θ*, fixation index *F_ST_* and the absolute nucleotide divergence *d_xy_*were calculated using the R package Popgenome [73].

### Identification of horizontal transfers

Putative horizontal transfers were first identified by mapping the best assembled reference genome representatives from the *F. oxysporum* and *F. fujikuroi* species complexes respectively to the *F. xylarioides* arabica563 reference genome. These were *F. verticillioides* (accession AAIM02000000) and *F. oxysporum* f. sp. *lycopersici* (accession AAXH01000000) respectively. Specifically, we looked for regions which were present in *F. xylarioides* and *F. oxysporum*, but absent from *F. verticillioides*. The *F. oxysporum* and *F. verticillioides* genomes were mapped to arabica563 using Minimap2 with the parameters “-cx asm10 –secondary=no –cs” [74]. Align-ment similarity was calculated in contiguous 10 kb windows, with similarity in each window calculated per 100 bp with no slide using “bedtools map -c 7 -o mean”. Regions were identified as putative horizontal transfers if they were absent from or less similar with *F. verticillioides*, than with *F. oxysporum*. The absence/ lower similarity in *F. verticillioides* was double checked using BLAST, whereby each putative HTR was tested against the genomes sequenced in this study (Table 2) and the BLAST nr database. A confirmed BLAST hit was only described if the target sequence was at least 50% in length and over 70% identity compared with the query sequence. This resulted in five putative HTRs identified in the *F. xylarioides* arabica563 genome on contigs 3, 6, 10, 11 and 13. After discovering that HTR 1 matched a chromosomal subregion “supercontig 51”, a region on the *F. oxysporum* mobile pathogenic chromosome which is enriched in effector genes, a BLAST match of each *F. oxysporum* chromosomal subregion returned an additional HTR in the robusta genomes (HTR 5).

Second, the phylogenetic branch distance for genes both inside and in 50kb regions flanking the putative horizontal transfers was calculated for arabica563 and *F. oxysporum* and *F. verticillioides* respectively. Gene presence in the *F. fujikuroi* complex species was determined using the orthogroup results from Orthofinder (*F. fujikuroi, F. proliferatum, F. verticillioides* and *F. anthophylum*, see Table S2). For those genes present in various *F. fujikuroi* complex species, phylogenetic branch distance was calculated using cophenetic in ape 5.0 in R [75]. Phylogenetic branch distance was also calculated between arabica563 and *F. oxysporum*. We only included as putative horizontal transfers those regions in which genes shared a higher percent identity between arabica563 and *F. oxysporum*, than arabica563 and the described *F. fujikuroi* complex species; this was four HTRs in the *F. xylarioides* arabica563 genomes on contigs 3, 6, 10 and 13, and one HTR in the *F. xylarioides* robusta genomes.

Non-vertical inheritance of certain CAZyme families was inferred using the gene trees from OrthoFinder for all CAZymes in the *F. xylarioides* reference genome. Phylogenetic distance between all clades was calculated using cophenetic in ape 5.0 in R [75].

*Starships* were identified following the parameters described in [19]. Briefly, regions absent from at least one genome were identified by aligning them to the reference genome using minimap2 and the parameters “-cx asm10” for a predicted distance of 10% divergence and “– secondary=no” to remove secondary alignments [74]. Bedtools was used to filter for contigs larger than 200kb, to remove repetitive regions and to create a non-redundant list of regions [76]. *Starship*-related genes were identified by their InterPro annotations. Starships were only verified where absent regions contained a “captain” gene (DUF3435) near their edge. The custom script used is available upon request.

### Infection assays and controls

The *F. xylarioides* strains IMI 389563 and IMI 375908 (“arabica563” and “arabica908”) from the arabica host-specific coffee wilt disease population from the CABI-IMI culture collection (CABI, Egham, UK) were used. Strains were grown on synthetic low nutrient agar at 25*^o^*C, with the spores verified after five days using lactophenol cotton blue staining under a compound microscope following [77]. The cultures were harvested with 1 drop of Tween20 and in 10ml sterile water. Two plates were combined to make each fungal inoculum before conidial concentrations were measured with a haemocytometer and adjusted to 10^6^ spores/ ml. A third control inoculum was made in the same way using sterile distilled water added to fresh synthetic low nutrient agar that had not been inoculated. Each inoculum type had four technical replicates.

*Coffea arabica* plants were purchased online from Gardeners Dream (https://www.gardenersdream.co.uk/) and left for 12 weeks to acclimate and grow in a controlled growth room at 25°C with a 12:12 hour light:dark cycle with 120 µmol/m2/s wavelength, and 50%:65% day:night humidity.

We compared gene expression of *F. xylarioides* strains infecting coffee plants and grown *in axenic* culture, as well as between the inoculated and control-inoculated coffee plants. Plants were infected with 10*µ*M through stem wounding [36], using a sterile needle at the first green (i.e. non-woody) internode. Each inoculum type (arabica563, arabica908, control inoculum) was used to inoculate four plants, making up 16 replicates in total per inoculum type. Plants were grouped by inoculum type and sealed in propagator trays with parafilm to make four trays containing four plants for each inoculum type. Tray location in the growth chamber was randomised and the plants were grown under shade at 25°C with a 12:12 hour light:dark cycle in a growth room until at least 50% of infected plants showed wilting, yellowing or early leaf senescence as symptoms of infection. The plants were harvested 98 days after inoculation.

Axenic cultures of arabica563 and arabica908 isolates were grown in GYM broth at 25°C with shaking at 175 RPM for seven days. As with the *in planta* inocula, the *in axenic* culture samples included four biological replicates (arabica563 1-4 and arabica908 1-4).

### RNA extraction, sequencing and quality control

All aboveground plant material was collected and flash-frozen in liquid nitrogen. The biological replicates were ground separately in liquid nitrogen with PVPP (polyvinyl polypyrrolidone) in a mortar and pestle. Approximately 5g of each biological replicate was pooled in a 1.5 ml flash-frozen polypropylene tube before vortexing and re-freezing.

Total RNA was extracted using the standard protocol of the RNeasy Mini kit (Qiagen, Germany). In total, 20 samples were extracted with four technical replicates for each treatment: *in planta* arabica563-infected coffee plants; *in axenic* culture arabica563; *in planta* arabica908-infected coffee plants; *in axenic* culture arabica908; control (-inoculated with water) coffee plants. RNA was quantified using a NanoDrop 2000 (Thermofisher), with the RNA quality assessment (Qubit and Fragment Analyzer), library preparation and Next Generation Sequencing (60M paired-end reads per sample) performed by Genewiz/ Azenta (Frankfurt, Germany). Raw sequenced reads were quality- and adapter-trimmed, see supplementary methods.

### Differential expression analysis

Using the Funannotate gene annotations as the target transcriptomes, transcript quantification was performed three times in a genome-free way using Salmon v1.5.2 [78] on the mapped read libraries: *in axenic* culture arabica563 and *in planta* arabica563; *in axenic* culture arabica908 and *in planta* arabica908; infected *C. arabica* plants and control-inoculated *C. arabica* plants. The relationships within and between biological replicates for each dataset were visually examined using the “PtR” script in the Trinity toolkit [56, 79]. Each transcript count matrix was tested for differential expression using DESeq2 [80], which uses negative binomial generalized linear models to test for statistical significance. Differential expression was also calculated using edgeR [81], which gave similar results, and limma/voom [82] which handles biases in RNA-seq data differently [83] and was too conservative with the read numbers from the *in planta* samples for this experiment. The Benjamini-Hochberg method [84] was used to adjust P-values for multiple testing to control the false discovery rate (FDR) and strict significance thresholds were applied, with P-values *<*0.001 and *>*4-fold to define a set of differentially expressed genes. The top 20% differentially expressed genes were classified as the “most differentially expressed”: 272 genes in arabica563; and 177 in arabica908.

### Functional annotation analysis

#### Gene Ontology and InterProScan classifications

Arabica coffee gene annotations (accession: GCF 003713225.1) were assigned InterProScan and Gene Ontology classifications using “interproscan.sh -iprlookup -goterms”.

#### EffectorP

A new set of putative effector proteins were identified using EffectorP 3.0 [85] on secreted proteins which were significantly differentially expressed in both fungal strains (total differentially expressed secreted proteins: 224 in arabica908, 380 in arabica563). Only secreted proteins significantly up-regulated in both strains were described as effector-like.

#### Carbohydrate-active enzymes

Carbohydrate Active EnZymes (CAZy) enzymes are grouped by family in the CAZy database[86]. CAZyme-encoding differentially expressed genes were identified in two ways, by comparing the proportion of differentially expressed genes to the genome and taking those where *>*50% of the genes were significantly differentially expressed in both strains, and by identifying enriched gene ontology terms with a cell-wall or pectin-degrading enzymatic function.

## Data analysis

All analyses were performed in R using R Statistical Software (v4.2.1; R Core Team 2022) [87] and in Linux run on the Imperial College Research Computing Service HPC facility (DOI: 10.14469/hpc/2232). See supplementary methods for a full list of R packages used to make figures.

## Acknowledgments

We thank W. Gordon, J. Meyer, M. Rutherford, T. Kibani, G. Armstong, D. Dring, R. Hillocks, E. Khonga and R. Mehrotra, as well as other un-named collectors who originally isolated the fungal pathogens and deposited them in the CABI-IMI collection, without which none of this work would have been possible. With thanks to the Imperial College Stevenson Fund for sponsoring L. D. Peck’s research placement in Genetique, Ecologie et Evolution at Université Paris Saclay, and to “i Focus & Write” and Barralab for endless support. With special thanks to A. Snirc for sequencing genomes, J. Vernadet for help with variant-calling, and S. Gurr and M. Fisher for invaluable thesis comments.

## Supporting information

### Materials and Methods

#### Strain details

All strains were from the CABI culture collection (Egham, UK), with IMI 507113 transferred from the Belgian coordinated collection of microorganisms from the Université Catholique de Louvain (Louvain-la-Neuve, Belgium) (under MUCL 47066/ MNHN 709) and IMI 507035 and 507038 transferred from the Agricultural Research Service culture collection (Illinois, USA) (with 507035 under NRRL 25804/ BBA 62721/ CBS 749.49; and 507038 under NRRL 37019/ FRC L96/ BBA 62458).

Each *F. xylarioides* MEGAHIT assembly from this study, as well as those assembled previously [26] and the publicly available genomes robusta925 [89] and robustaL0394 [90], were mapped to the reference genome assembly.

Single nucleotide polymorphisms (SNPs) were called against the arabica563 reference genome for each set of *F. xylarioides* reads. Reads for robusta925 were downloaded from the Sequence Read Archive (accession PRJNA508603, [89]) and robustaL0394 were shared by [90].

#### Illumina genome assembly

Low quality bases (Phred score *<*20) and adapters (stringency 4) were removed using TrimGalore 0.6.0 by Cutadapt (https://github.com/FelixKrueger/TrimGalore, [91]. Overlapping paired-end reads were stitched together using FLASH 1.2.11 [92] and *de novo* assembled with MEGAHIT 1.2.9 [93, 94]. Assembly metrics including BUSCO scores were computed using QUAST 5.2.0 (Table 2) [95].

#### Long-read reference genome sequencing and assembly

High-quality high molecular weight genomic DNA was extracted from 1-2 g of mycelium ground in liquid nitrogen using a lysis buffer containing sodium metabisulfite, 2M Tris, 500 mM EDTA, 5M NaCl, and 5% Sarcosyl. The gDNA was extracted from the cell lysate using Chloroform, and precipitated using Isopropanol and 70% ethanol, following the protocol of [51]. The gDNA was eluted in 500*µ*l TE buffer and incubated overnight at 4*^o^*C. RNA contaminants were removed using RNAse A (10mg/ ml) and gDNA was purified using Agencourt AMPure XP beads (ThermoFisher).

#### Orthologous gene groups

Orthogroups were inferred using the OrthoFinder algorithm [96] with unrooted gene trees built for each orthogroup using DendroBLAST [97]. Trees were resolved using the OrthoFinder hybrid species-overlap/ duplication-loss coalescent model [96], and drawn in ggtree [98].

#### RNA quality control

Raw sequenced reads were quality- and adapter-trimmed using Trimmomatic v0.39 (parameters: ILLUMINACLIP:$TRIMMOMATIC DIR/adapters/TruSeq3-PE.fa:2:30:10 SLIDINGWINDOW:4:5 LEADING:5 TRAILING:5 MINLEN:25) [99]. Reads were quality-checked with FASTQC v0.11.2 [100] and retained where both pairs were trimmed. To remove ribosomal RNA sequences, reads were mapped to the SILVA rRNA database using BBTools “bbmap” with the parameter “outu=filtered R*.fq.gz” to keep only the unmapped reads. Finally, the reads were repaired using BBTools “bbmap” “repair.sh” to re-pair reads that became disordered or had their pair eliminated. These QC steps removed 61 GB of data. Before using the data to quantify gene expression, reads were mapped using “bbmap.sh” to the gene annotations of arabica563 and arabica908 and from *C. arabica* (GCA 003713225.1, [101]). These mapped read libraries were used for all next steps.

#### Data presentation

The following R packages were used to make figures, all using the latest versions: tidyverse [102], cowplot [103], ggrepel [104], wes anderson [105], pheatmap [106], ggtree [98], RIdeogram [107], circlize [108], ape [75], viridis [109], pals [110], ggbreak [111], pafr [112], treeio [113], ggfortify [114].

### Supplementary Results

#### A high-quality genome assembly for the *Fusarium xylarioides* arabica host specialist

Assembly of Nanopore long-read data for *F. xylarioides* IMI 389563 (“arabica563”) generated its first high-quality reference genome with 21 contigs, including 12 contigs over 1 Mb, and four contigs over 100 kb. Telomeric repeats were identified on both ends of three chromosomes, consistent with these representing complete assembled chromosomes, and at just one end of seven contigs, probably representing incomplete chromosomes (Figure S3B). The assembly length was 57 Mb (with 55 Mb in the 12 longest contigs) with an N50 greater than 5 Mb (Table 2). The RagTag assemblies ordered and oriented to the reference genome for the remaining *F. xylarioides* strains allowed the identification of various large arabica-specific regions over 0.7 Mb in contigs 6, 7, 8, 12 and 13, compared to robusta-specific genomes. The regions in contigs 7, 12, and 13 were absent from all robusta- and coffea-specific genomes.

#### Differential gene expression

Among the expressed genes and relative to their relevant controls, an average of 2164 were differentially expressed across the arabica563 samples; 1713 across the arabica908 samples; and 233 across the coffee plant control samples. In the fungal samples, over half of the differentially expressed genes were upregulated *in vivo* infection samples relative to controls, whilst in the coffee plant samples 80% of the differentially expressed genes (116 in the plants infected with arabica563; 231 with arabica908) were up-regulated in the infected samples. Genes which were differentially expressed in both *F. xylarioides* arabica strains belong to nearly 700 orthologous groups. The differentially expressed coffee genes are described in Table S11.

### Supplementary Figures and Tables

**Figure S1:**
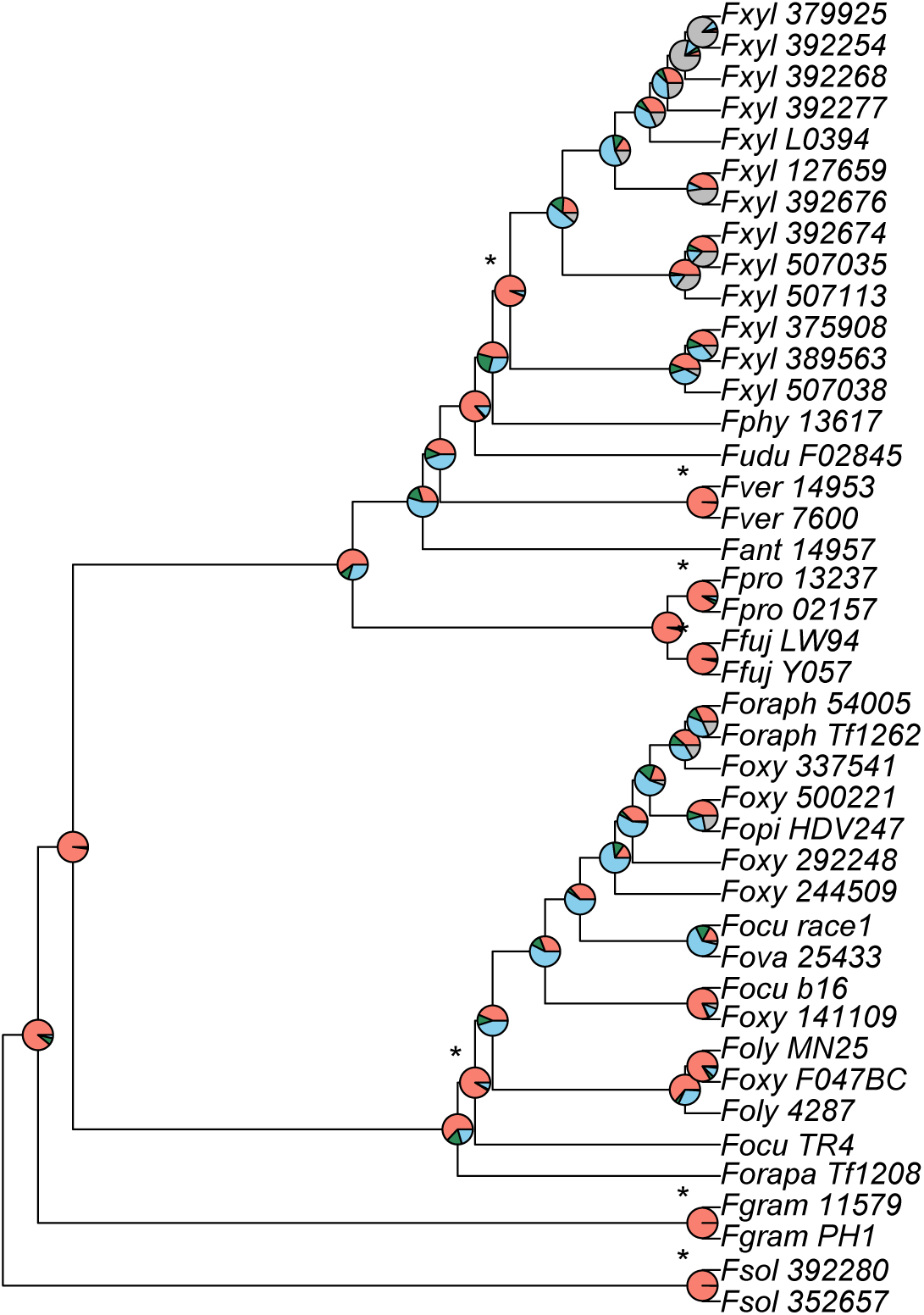
Phylogenetic relationships between *Fusarium* genomes reconstructed from 1685 single copy orthogroups support monophyly across all identified species clades, with little consistent support for alternate topologies. Species tree built in ASTRAL. Full strain accession numbers are in Table S2. Pie chart colours: pink = proportion of genes(orthogroups) recovering the depicted node; dark green = the proportion of genes recovering the second most common topology; light blue = the proportion of genes recovering all other topologies; grey = unresolved topology.

**Figure S2:**
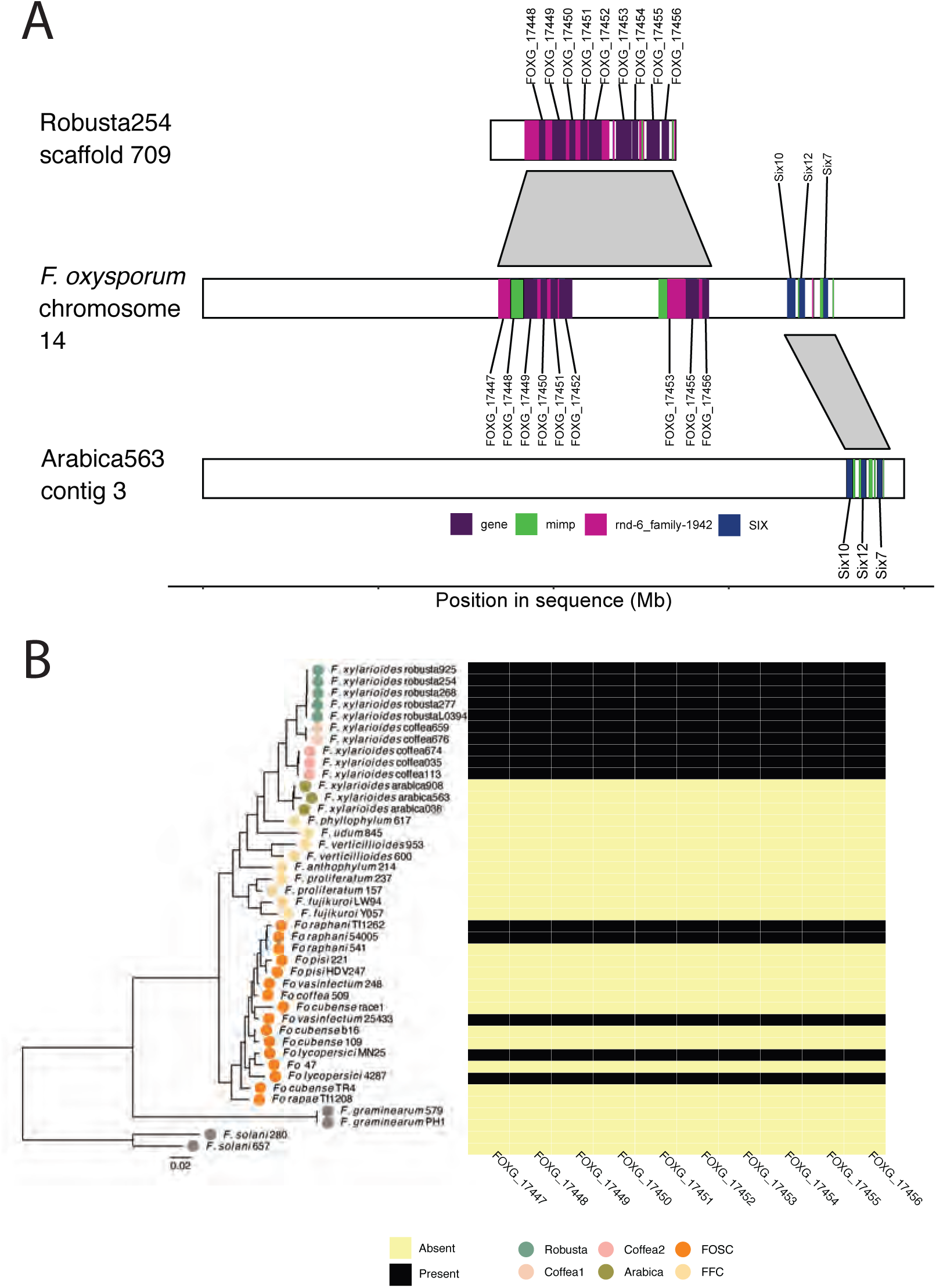
A large, gene-rich region is shared between *Fusarium oxysporum* f. sp. *lycopersici* and *Fusarium xylarioides* arabica and robusta genomes. **A** Separate regions of a highly similar region in *Fusarium oxysporum* f. sp. *lycopersici* chromosome 14 (NC 030999.1, middle plot) match to *F. xylarioides* robusta scaffold 709 (top plot) and *F. xylarioides* arabica contig 3 (bottom plot). Genes shared by the two species are shaded according to the legend and labelled with the Fusarium oxysporum gene locus tag. Repeats of interest are shaded according to the legend. Each shared gene in this region also shares an orthologous group with *Fusarium oxysporum*. Drawn to scale. **B** Heatmap showing presence of orthologous groups found in the region found in *F. xylarioides* robusta from (A), compared with *Fusarium* phylogeny. Present genes are shaded in dark and all share *>*96% identity. The phylogenetic species tree was created following Figure 2. Full strain details in Tables S2 and 2.

**Figure S3:**
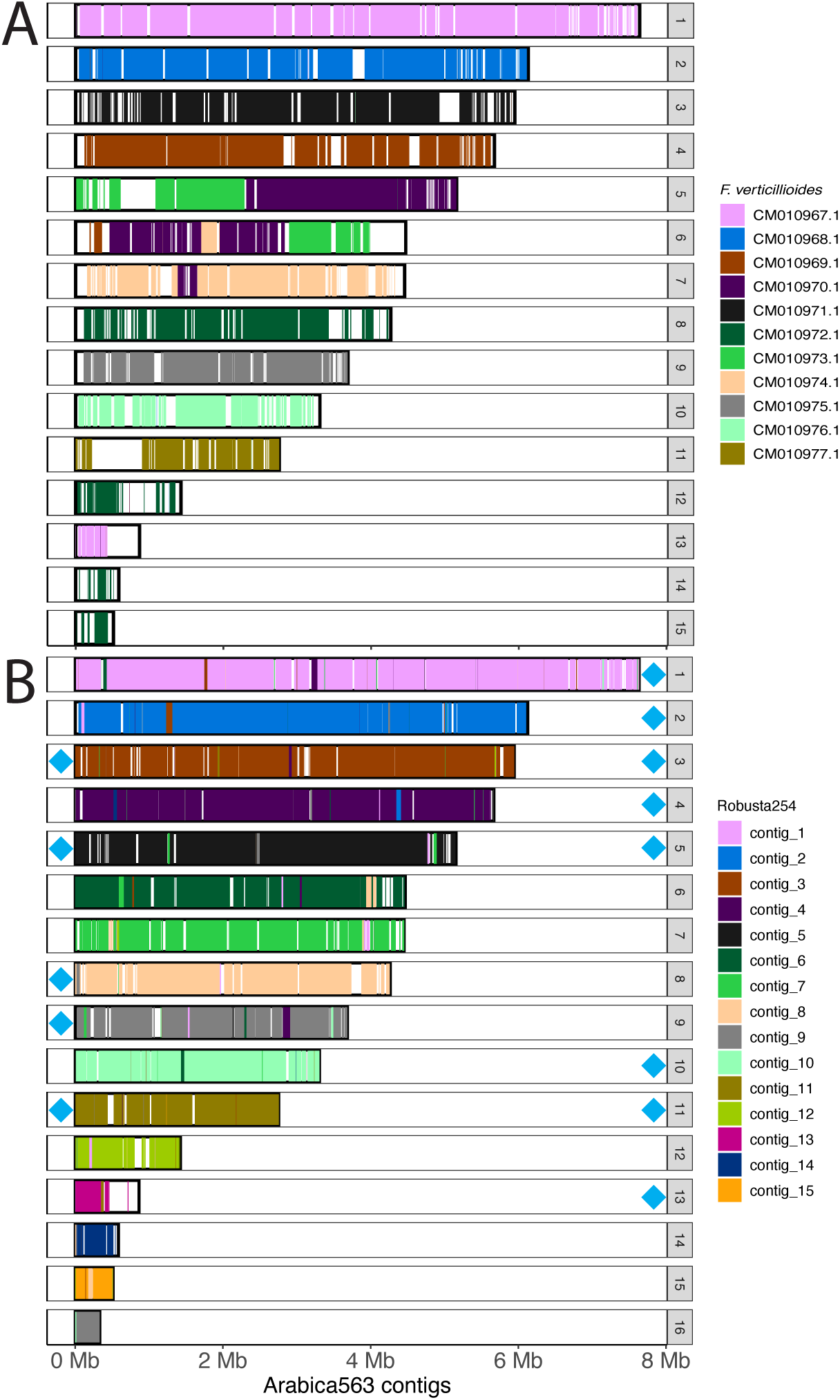
Whole-genome alignments (A) *Fusarium verticillioides* chromosomal assembly mapped against the *Fusarium xylarioides* arabica563 contig assembly and (B) *Fusarium xylarioides* robusta254 mapped against the *Fusarium xylarioides* arabica563 contig assembly. **A** Each bar represents one of the 16 arabica563 contigs *>*100 kb in length that are also present in *Fusarium verticillioides*, with contig size and position on the x axis. Arabica563 contig numbers are annotated in-line with the bars. Shaded regions indicate the presence of syntenic *Fusarium verticillioides* chromosomes, the colour of which identifies the specific mapped chromosome. Unshaded regions indicate those found only in arabica563, and arabica563 contig 16 is absent from *Fusarium verticillioides*. *Fusarium verticillioides* chromosomes CM010967.1-CM010977.1 correspond to 1-11 in the genome accession GCA_000149555.1. **B** Each bar represents one of the 16 *Fusarium xylarioides* arabica563 contigs that are also present in *Fusarium xylarioides* robusta254, with contig size and position on the x axis. Shaded regions indicate the presence of syntenic *Fusarium xylarioides* robusta254 contigs, the colour of which identifies the specific mapped contig from *Fusarium xylarioides* robusta254. Unshaded regions indicate contigs found only in *Fusarium xylarioides* arabica563 that are absent in *Fusarium xylarioides* robusta254. Telomeric repeats are annotated as blue diamonds on their corresponding scaffolds. Contigs *<*1 kb not shown.

**Figure S4:**
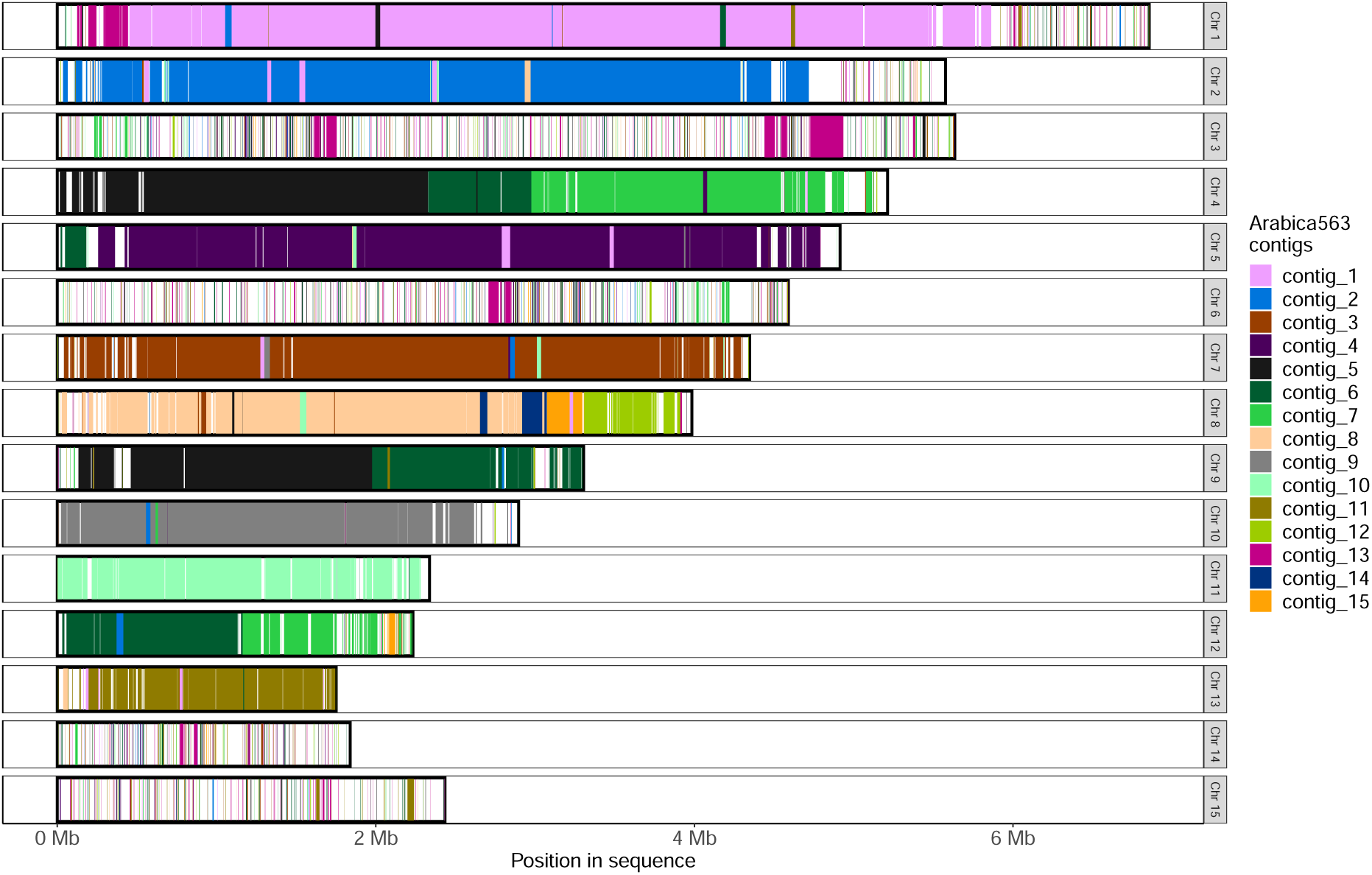
Whole-genome alignment of the *Fusarium xylarioides* arabica563 contig assembly mapped against *Fusarium oxysporum* f. sp. *lycopersici* chromosomal assembly. Each bar represents one of the 15 *Fusarium oxysporum* f. sp. *lycopersici* chromosomes that are also present in *Fusarium xylarioides* arabica563, with contig size and position on the x axis. Arabica563 contig numbers are annotated in-line with the bars. Shaded regions indicate the presence of syntenic Arabica563 contigs, the colour of which identifies the specific mapped contig. Unshaded regions indicate those found only in *Fusarium oxysporum* f. sp. *lycopersici*. *Fusarium oxysporum* f. sp. *lycopersici* chromosomes 1-15 correspond to NC_030986-NC_031000 in the genome accession GCA_000149555.1.

**Figure S5:**
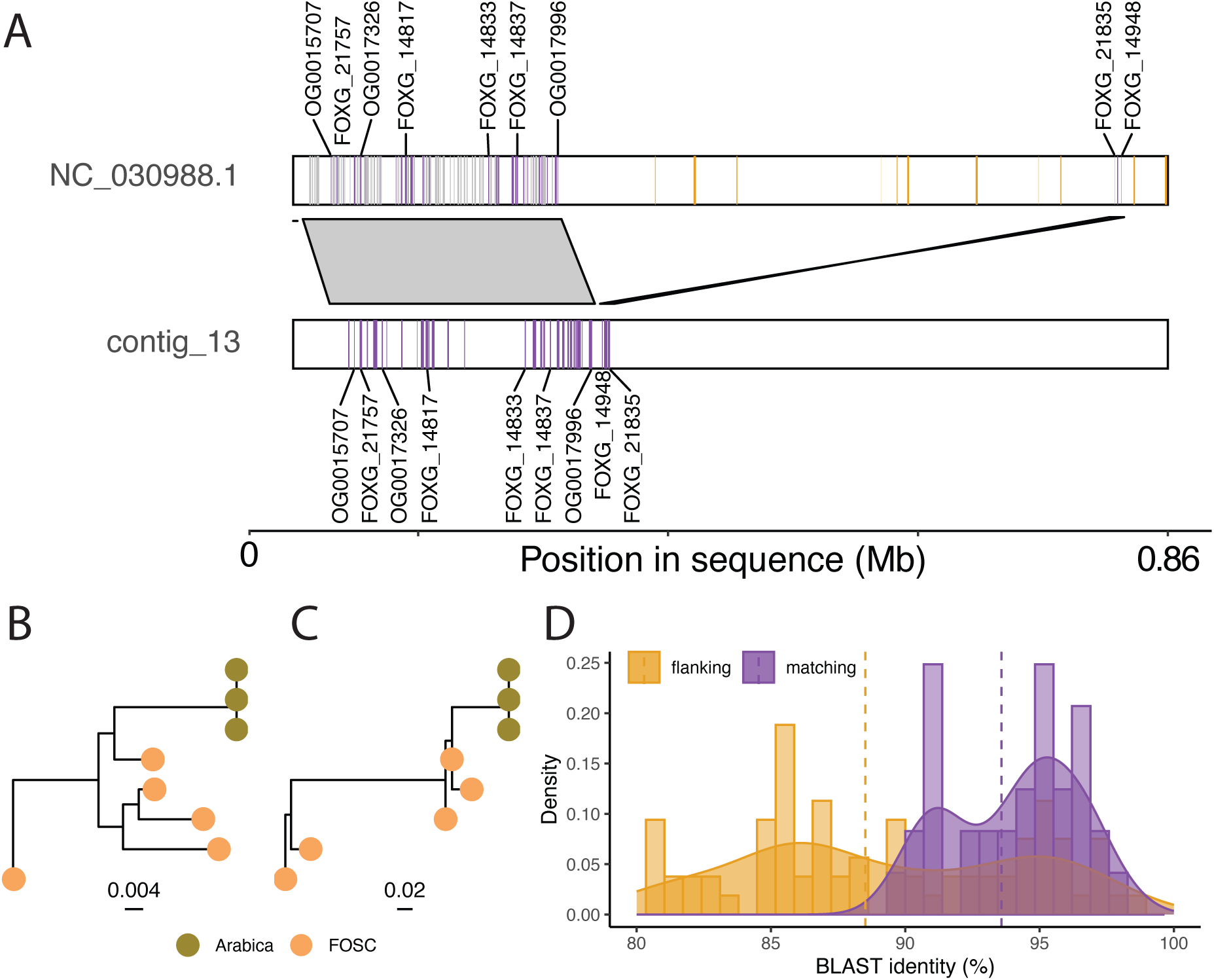
Putative horizontal transfers between *Fusarium oxysporum* and *Fusarium xylarioides*. **A** Contig 13 in *Fusarium xylarioides* arabica563 (544 kb - 739 kb, bottom plot) matches two regions in *Fusarium oxysporum* f. sp. *lycopersici* chromosome 3 (NC 030988, 4.7 Mb - 5.6 Mb, top plot) in two pieces, large (200kb) and small (20kb). Highly similar genes shared by the two species are shaded in purple in top and bottom plots, a subset are labelled by *Fusarium oxysporum* gene locus tag, and non-matching genes found in the shared region are shaded grey. Genes flanking the shared region in *Fusarium oxysporum* are shaded yellow, and those which are found in *Fusarium xylarioides* (BLAST length *>*80%) have a lower sequence identity and are absent from contig 13. **B** A DendroBLAST rooted gene tree for the arabica563 gene P1J47 002169 (OG0015707) orthologous group, found in the shared region. **C** DendroBLAST gene tree for the arabica563 gene P1J47 003005 (OG0017996) orthologous group, found in the shared region. **D** The genes shared between the two species in the matching region have a higher BLAST sequence identity (mean 94%, dotted line) than those found in the flanking regions either side (mean 88%, dotted line). Figure drawn to scale.

**Figure S6:**
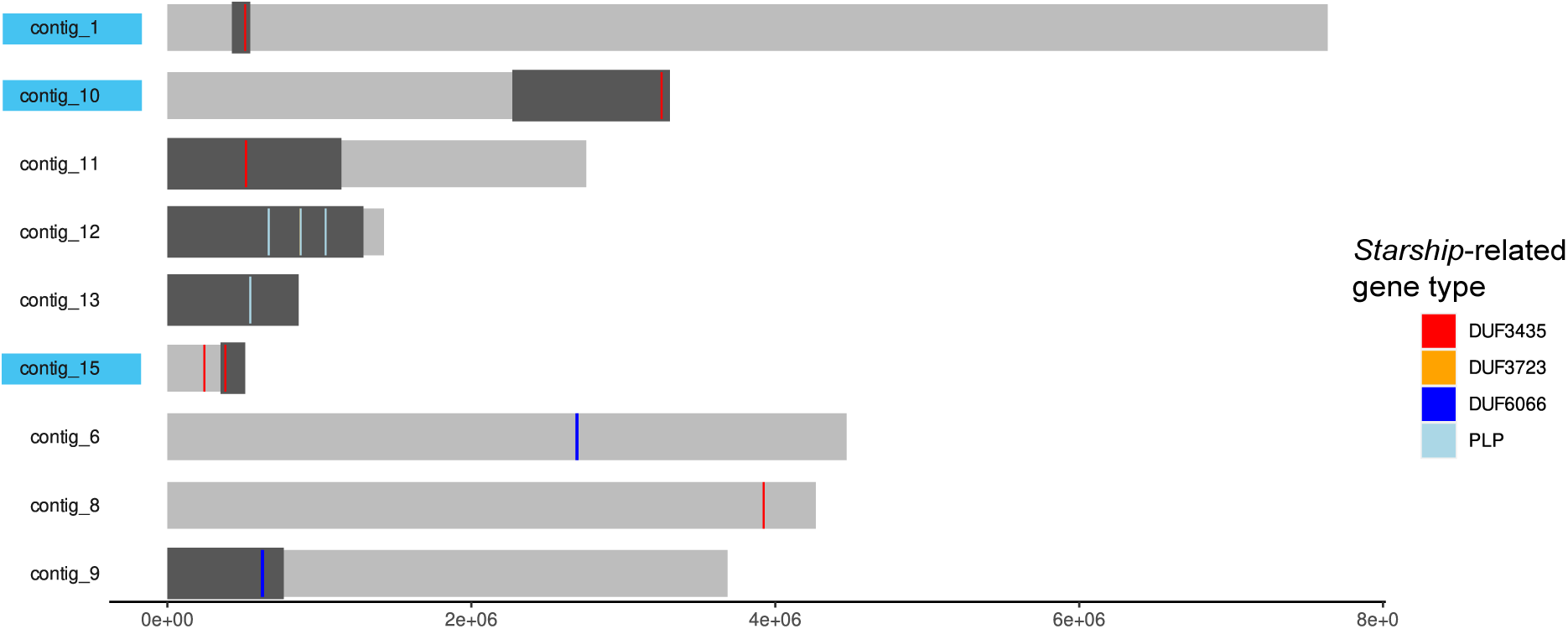
*Starships* in the arabica563 reference genome. Each region which is present in arabica563 but absent from at least one other clade (including *F. oxysporum* and members of the *F. fujikuroi* species complex) is shaded dark grey. Conserved *Starship* cargo genes which were identified in arabica563 are annotated and include “captains”, DUF3435, as well as *Starship*-related genes that captains often co-localise with and include DUF3723, DUF6066 and patalin-like phosphatases (PLPs). Contigs which contain a confirmed *Starship* are shaded with a blue box, namely those which contain a captain at the edge of the absent (shaded dark grey) region. Only contigs containing *Starship*-related genes are shown.

**Figure S7:**
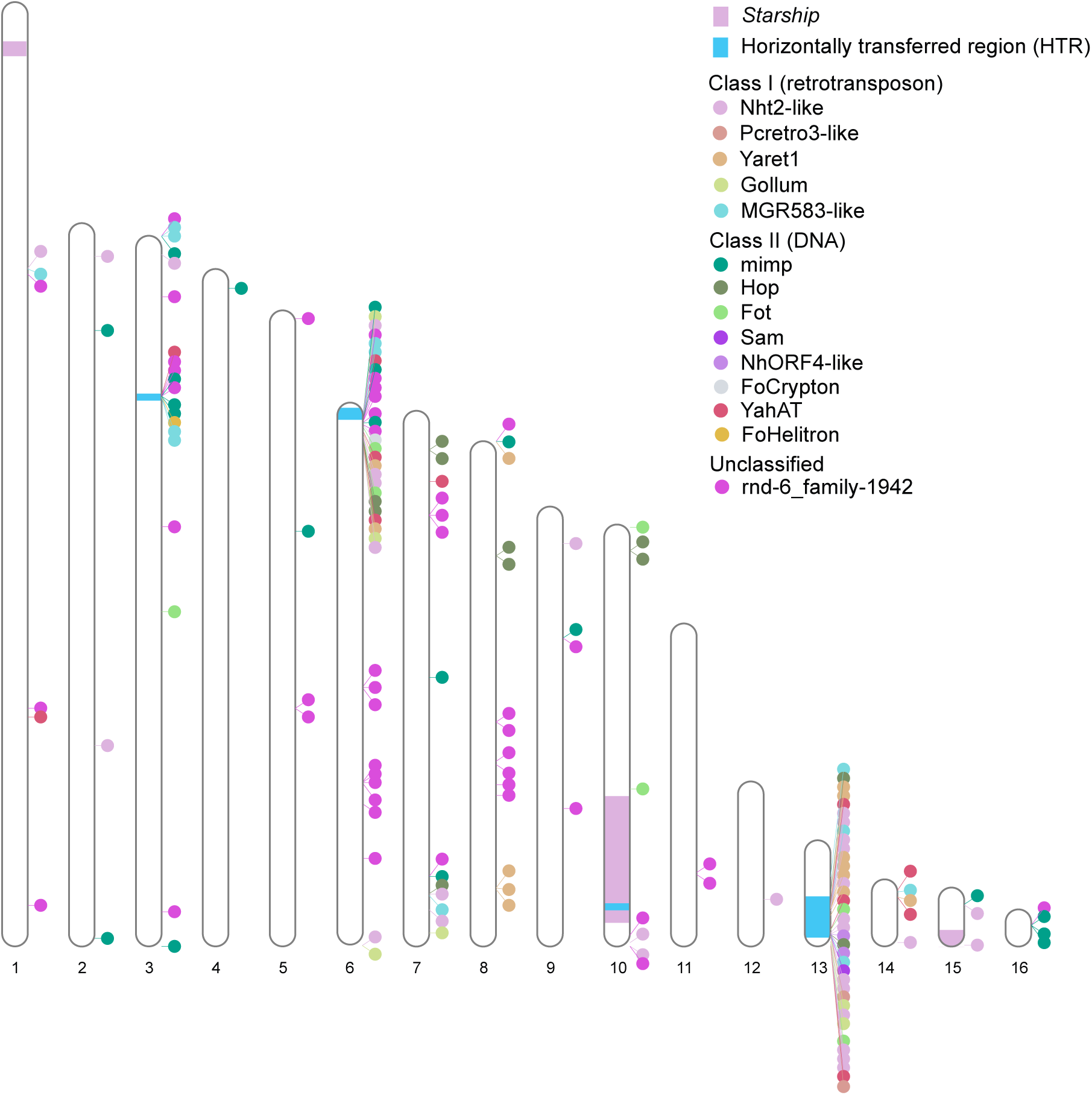
Whole-genome view of the loci of transposable elements in the *F. xylarioides* arabica563 reference genome. Transposable elements shown are those which were newly identified on the mobile and pathogenic chromosome of *F. oxysporum* f. sp. *lycopersici* by [30], as well as the unclassified family rnd-6 family-1942. Contigs *<* 100 kb are not shown.

**Table S1:**
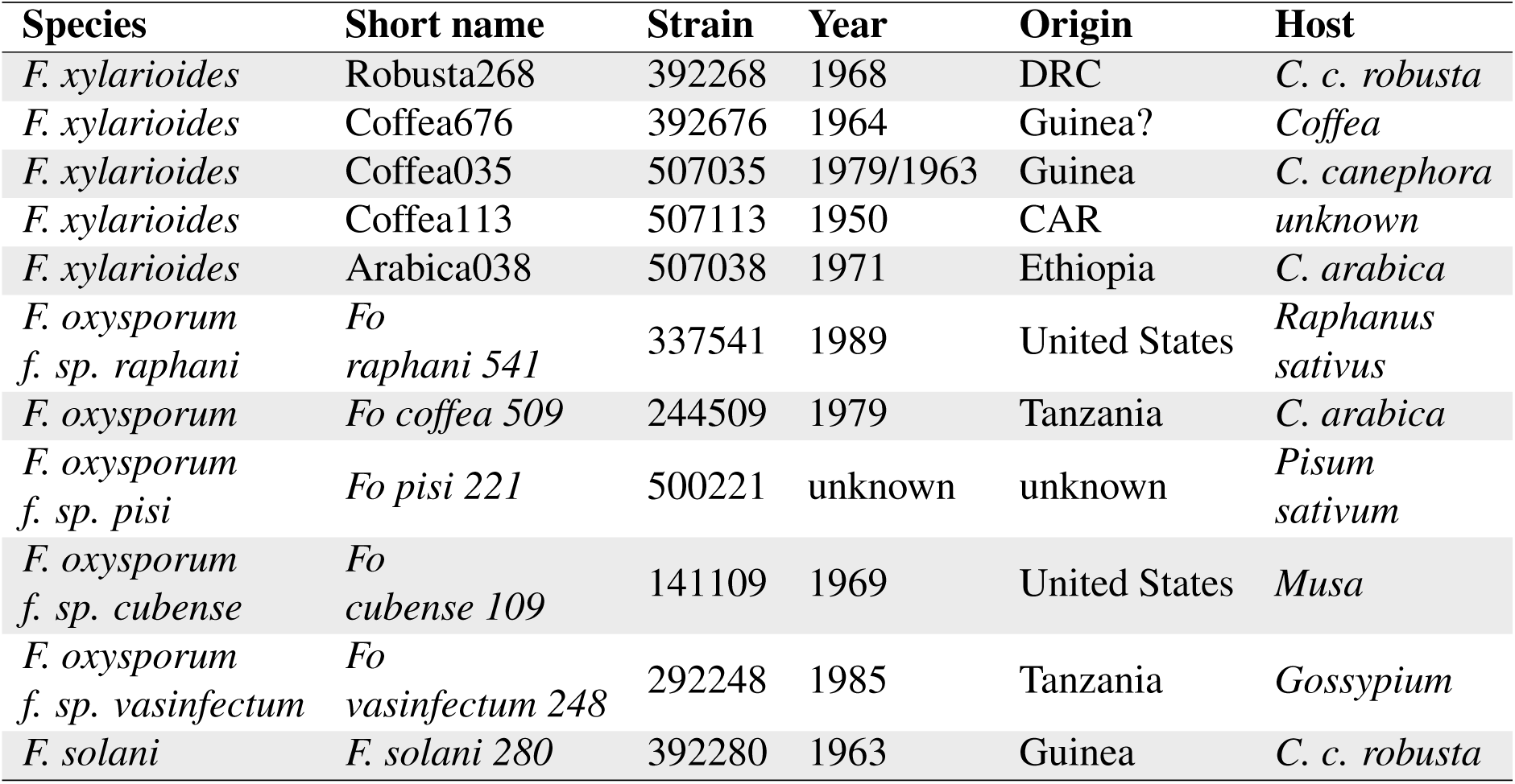
Bio-geographic details for strains sequenced in this study.

**Table S2:**
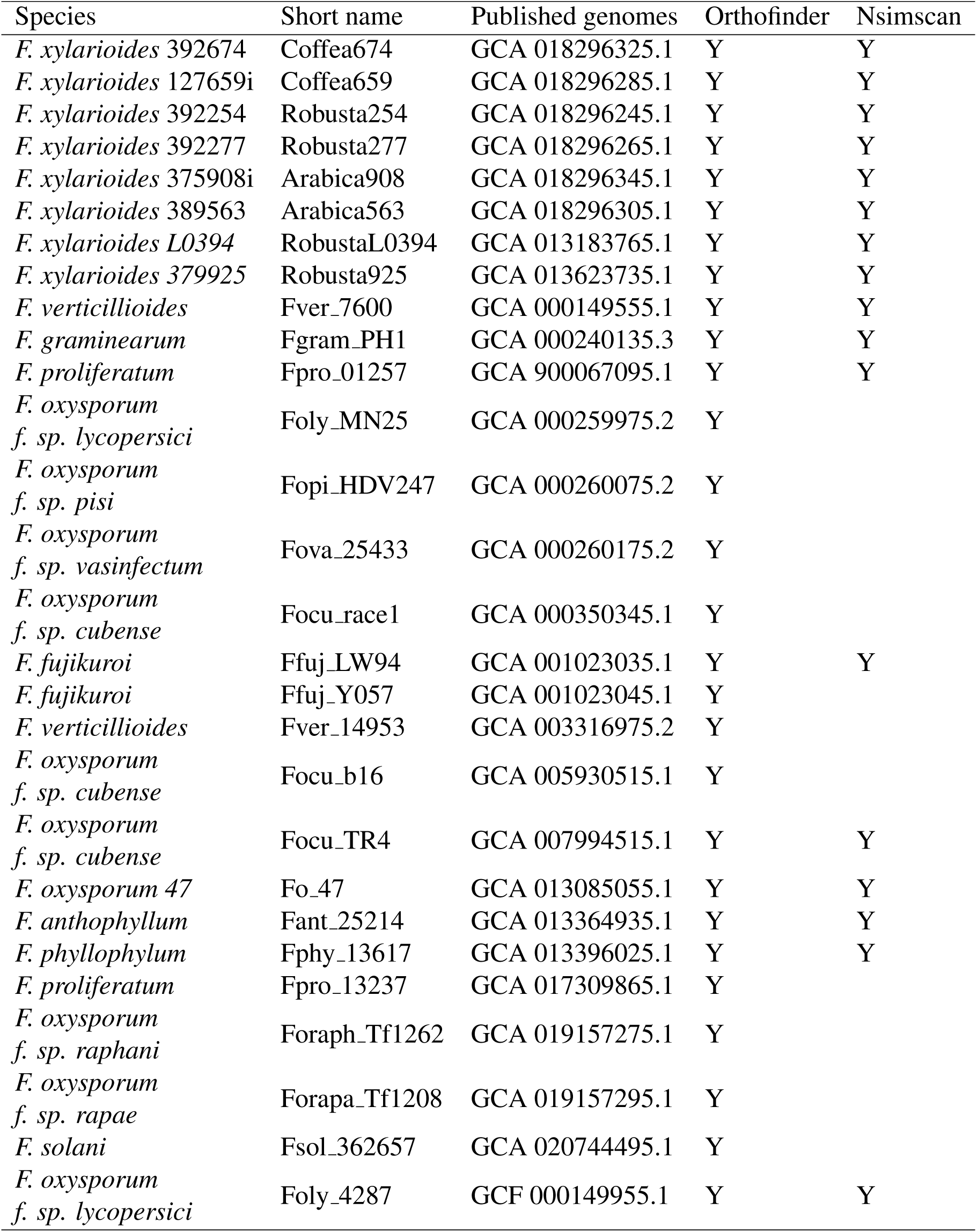
Published genomes analysed in this study. The columns describe: species, short species with strain number, accession number, whether the genome was used in OrthoFinder, whether the genome was used in Figure 2B for whole-genome similarity.

**Table S3:**
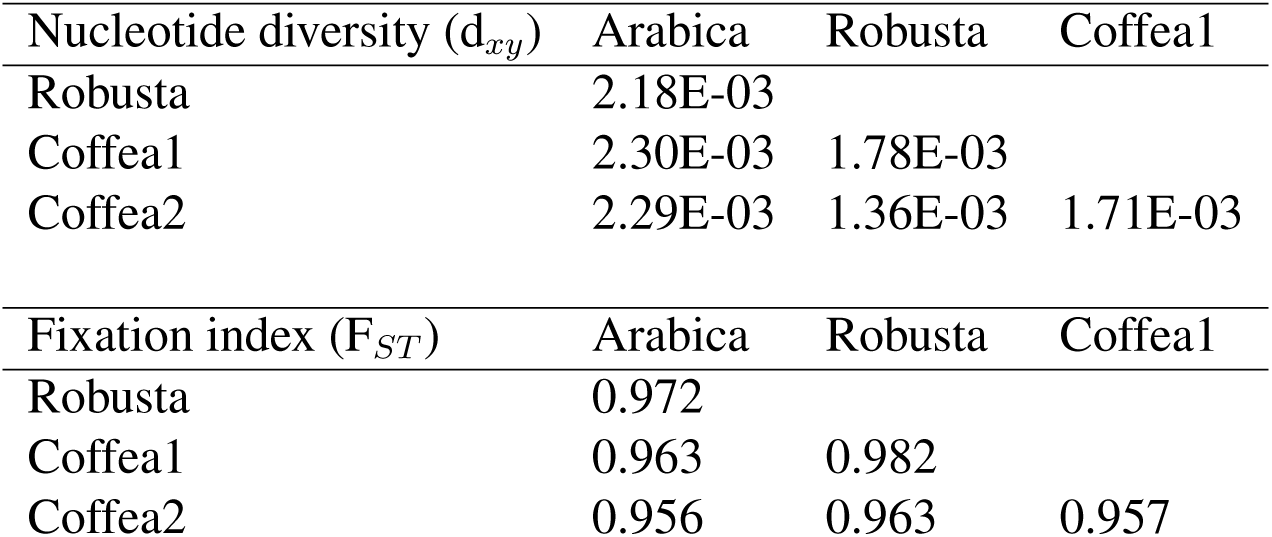
Population genetics statistics related to genetic differentiation (F*_ST_*, d*_xy_*) in four *Fusarium xylarioides* populations. Measured in 100 kb windows.

**Table S4:**
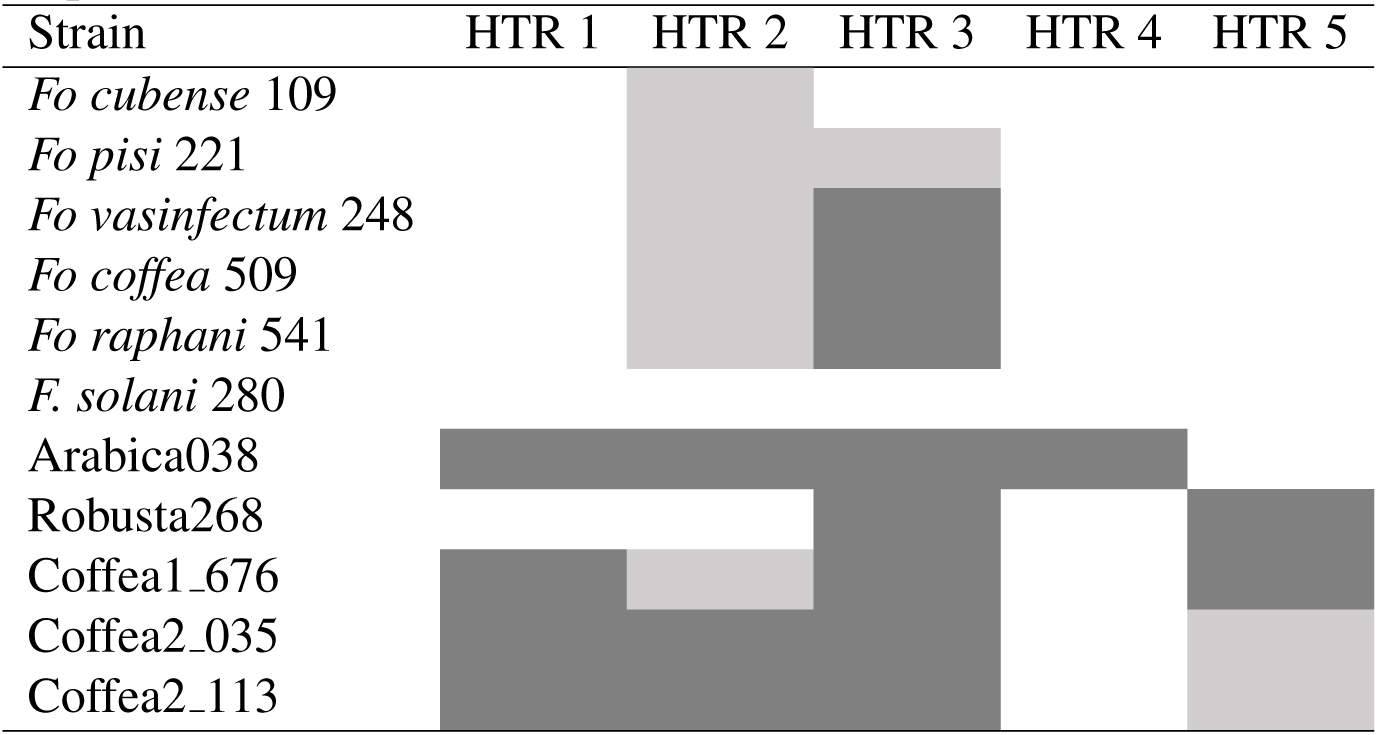
The presence of putative horizontally transferred regions (HTR) across *Fusarium* genomes sequenced in this study. Light shading indicates sequences with a BLAST hit that is *>*50% length and *>*70% identity compared with the arabica563 HTR sequence; dark shading indicates a BLAST hit that is *>*80% length and *>*80% identity. Abbreviations: *Fo*, *F. oxysporum* f. sp.

**Table S5:**
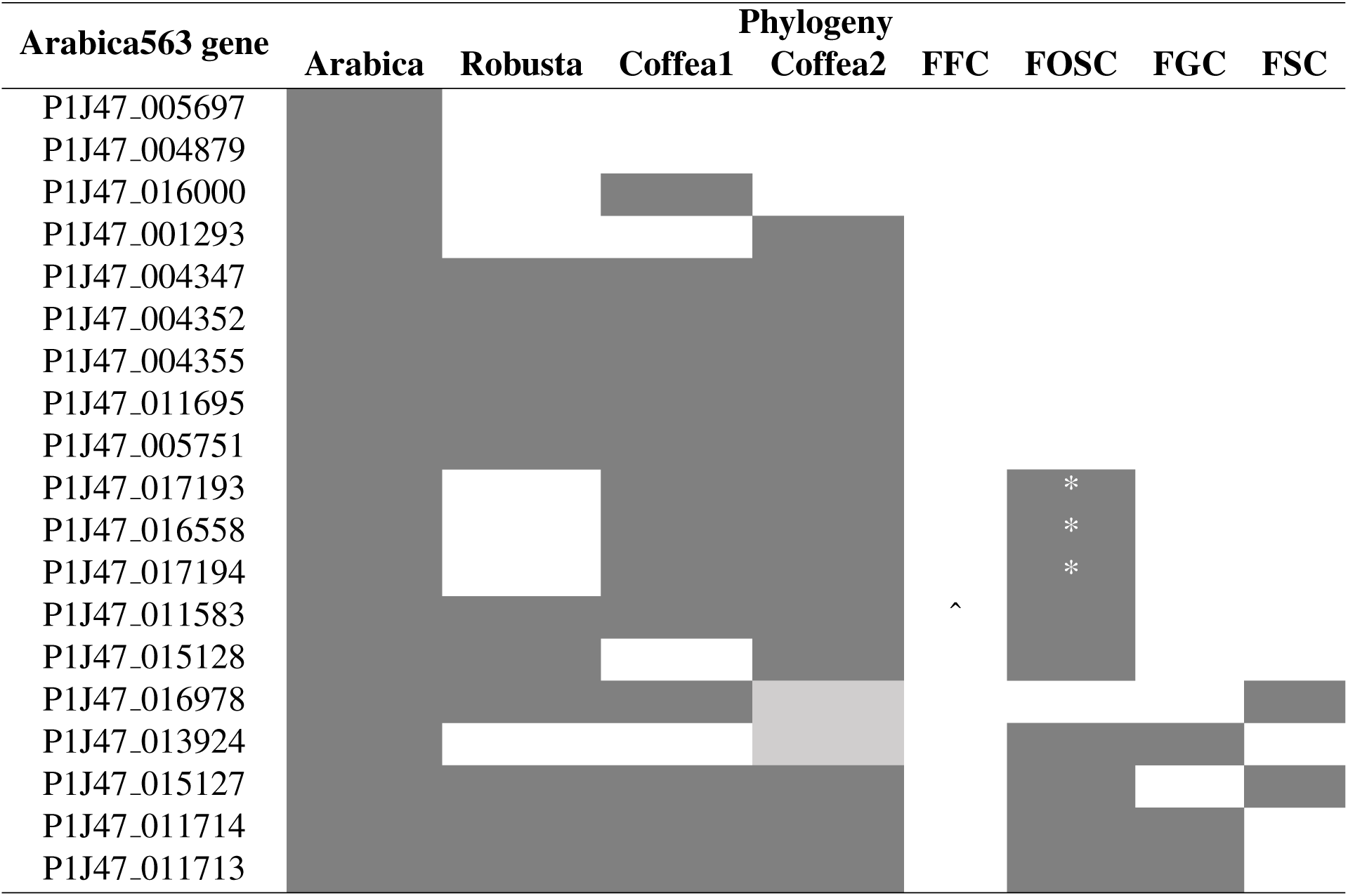
Nineteen up-regulated genes in both *Fusarium xylarioides* arabica strains are absent from the *Fusarium fujikuroi* complex (FFC) and differentially present across *Fusarium xylarioides*, the *Fusarium oxysporum* species complex (FOSC), the *Fusarium graminearum* species complex (FGC) and the *Fusarium solani* species complex (FSC). Species presence was determined using genealogies from OrthoFinder (see methods) and BLAST similarity. A caret indicates a genealogy present in *Fusarium udum*, the other vascular wilt in the *Fusarium fujikuroi* species complex. Partial shading indicates partial presence across the population. An asterisk represents genealogies which match *Fusarium oxysporum* effector proteins.

**Table S6:**
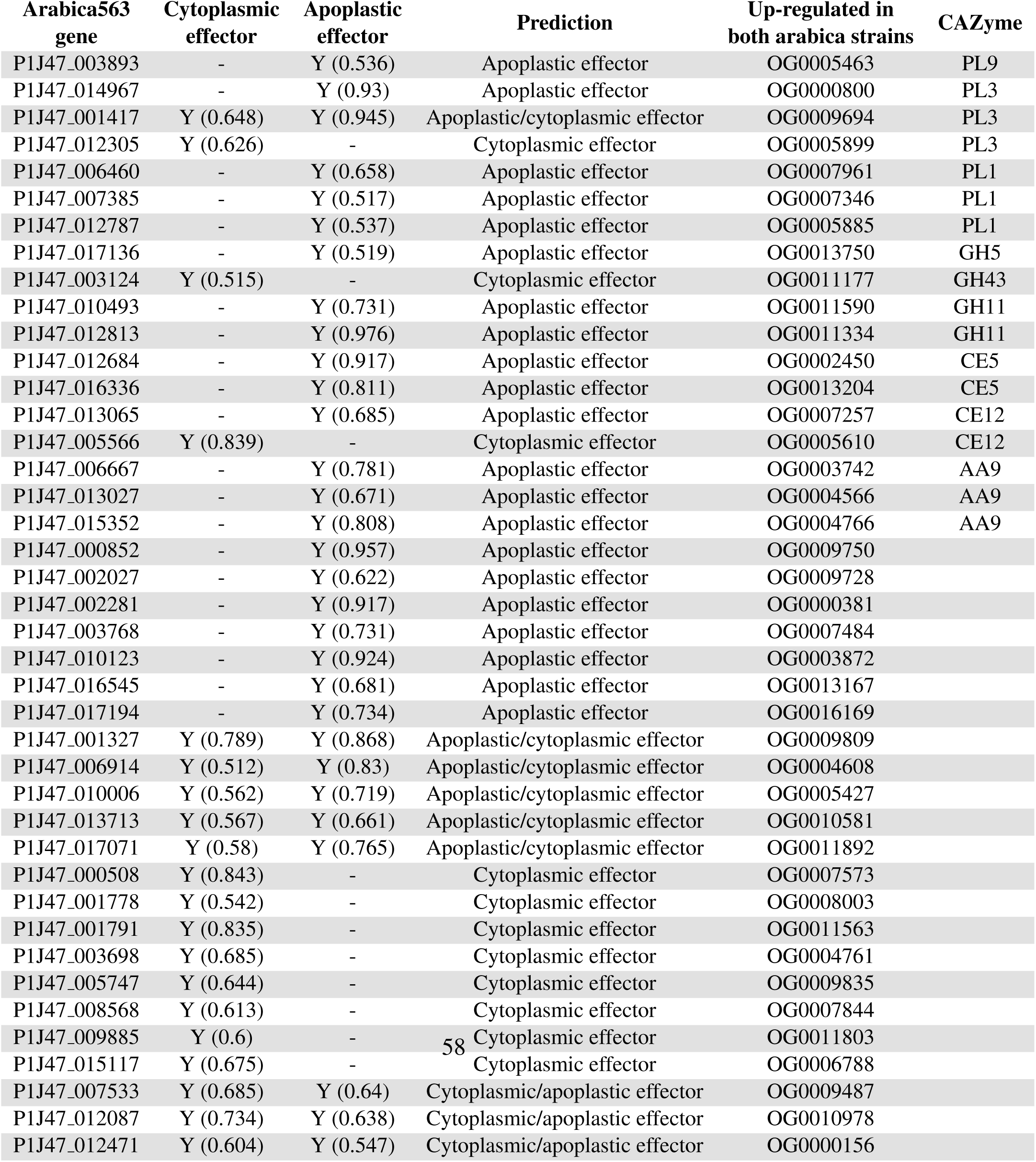
Predicted effectors in arabica563 genes by EffectorP [85]. Only those which were up-regulated in both *Fusarium xylarioides* arabica563 and arabica908 are shown. The numbers in brackets describe the certainty. The orthogroup is shown for those predicted effectors up-regulated in both arabica563 and arabica908, and those which are annotated as carbohydrate-active enzymes are described. P1J47 010006 encodes an Ecp2 effector protein and P1J47 007533 contains a LysM domain.

**Table S7:** Expression data for all putative effectors from [26] across the *Fusarium xylarioides* arabica563 samples. Each gene is described as up-regulated *in planta* (“coffee.up”), *in axenic* (“culture.up”) or not differentially expressed (“ns”).

| Arabica563 gene | log2FoldChange | False Discovery Rate | Predicted EffectorP? | Differentially expressed? | Putative effector |
| --- | --- | --- | --- | --- | --- |
| P1J47_011695-T1 | 13.4154519 | 7.18E-47 | FALSE | coffee.up | OG0014398 |
| P1J47_015569-T1 | 5.379337 | 3.84E-29 | FALSE | coffee.up | FOXG_14254 |
| P1J47_014967-T1 | 14.4073093 | 2.66E-16 | TRUE | coffee.up | pelA |
| P1J47_004459-T1 | 11.736764 | 4.71E-16 | TRUE | coffee.up | pelD |
| P1J47_002381-T1 | 2.5053746 | 1.88E-10 | FALSE | coffee.up | orx1 |
| P1J47_017001-T1 | 4.9013526 | 2.62E-09 | FALSE | coffee.up | OG0018569 |
| P1J47_007374-T1 | -4.8076726 | 2.45E-07 | FALSE | culture.up | OG0014367 |
| P1J47_007948-T1 | 2.3398743 | 2.93E-06 | FALSE | coffee.up | sge1 |
| P1J47_014428-T1 | 3.9418829 | 7.27E-05 | FALSE | coffee.up | catalase-peroxidase |
| P1J47_013555-T1 | 1.0310702 | 9.98E-04 | FALSE | ns | fow1 |
| P1J47_005049-T1 | -0.7238584 | 4.23E-03 | FALSE | ns | snf1 |
| P1J47_012661-T1 | -2.5494558 | 3.20E-02 | FALSE | ns | chlo_vacu |
| P1J47_013807-T1 | -2.0638231 | 5.75E-02 | FALSE | ns | OG0014828 |
| P1J47_001679-T1 | -2.871467 | 1.09E-01 | FALSE | ns | OG0013912 |
| P1J47_002427-T1 | 0.3631168 | 1.23E-01 | FALSE | ns | rho1.1 |
| P1J47_003388-T1 | -1.6549056 | 1.82E-01 | FALSE | ns | OG0013871 |
| P1J47_013972-T1 | -0.2587035 | 4.35E-01 | FALSE | ns | rho1.2 |
| P1J47_016733-T1 | -0.3777736 | 5.01E-01 | FALSE | ns | OG0013877 |
| P1J47_016313-T1 | 0.141086 | 8.53E-01 | FALSE | ns | fmk1 |
| P1J47_017133-T1 | 15.6417987 | 1.00E+00 | TRUE | ns | OG0013477 |

**Table S8:**
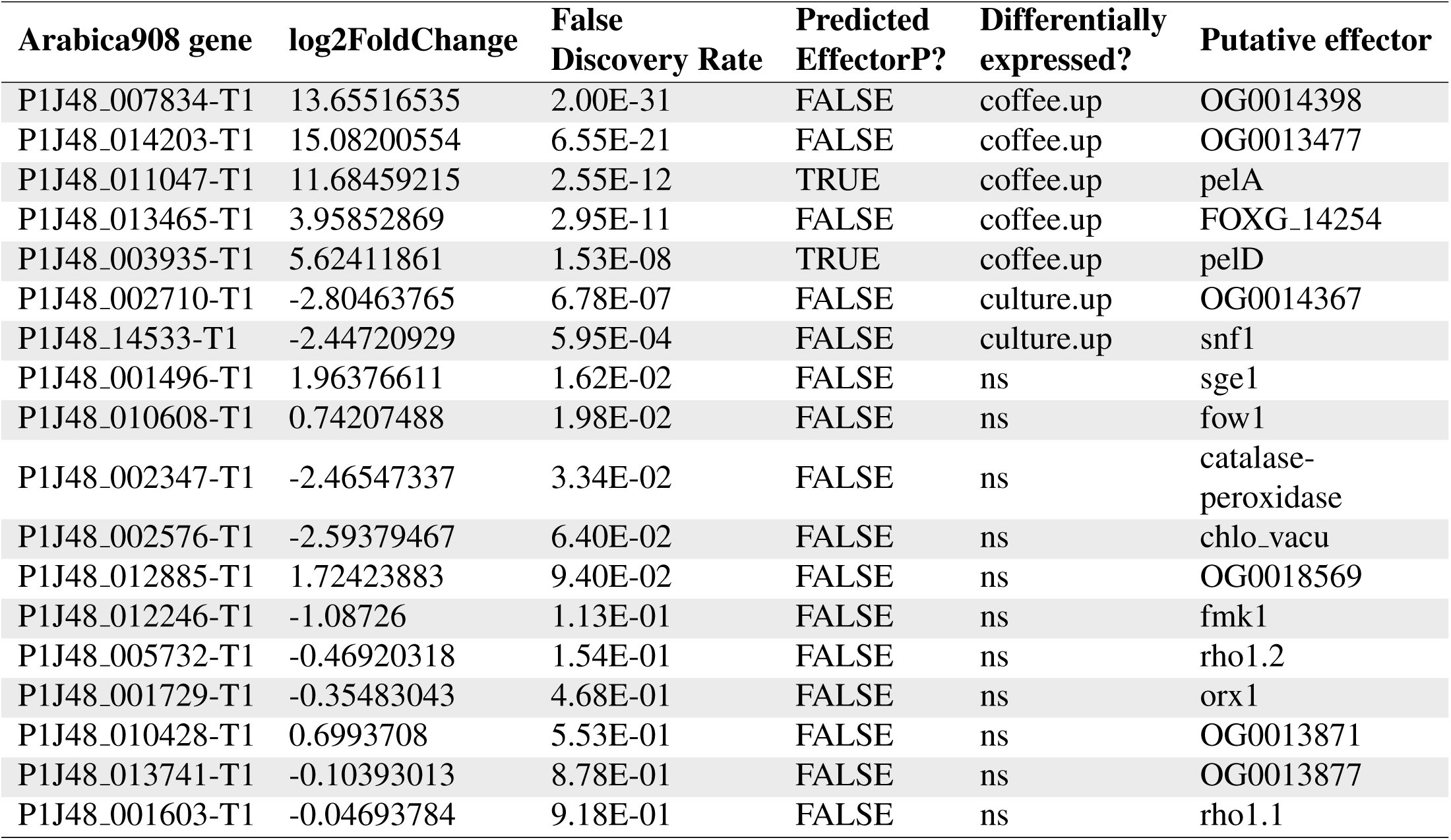
Expression data for all putative effectors from [26] across the *Fusarium xylarioides* arabica908 samples. Each gene is described as up-regulated *in planta* (“coffee.up”), *in axenic* (“culture.up”) or not differentially expressed (“ns”).

**Table S9:**
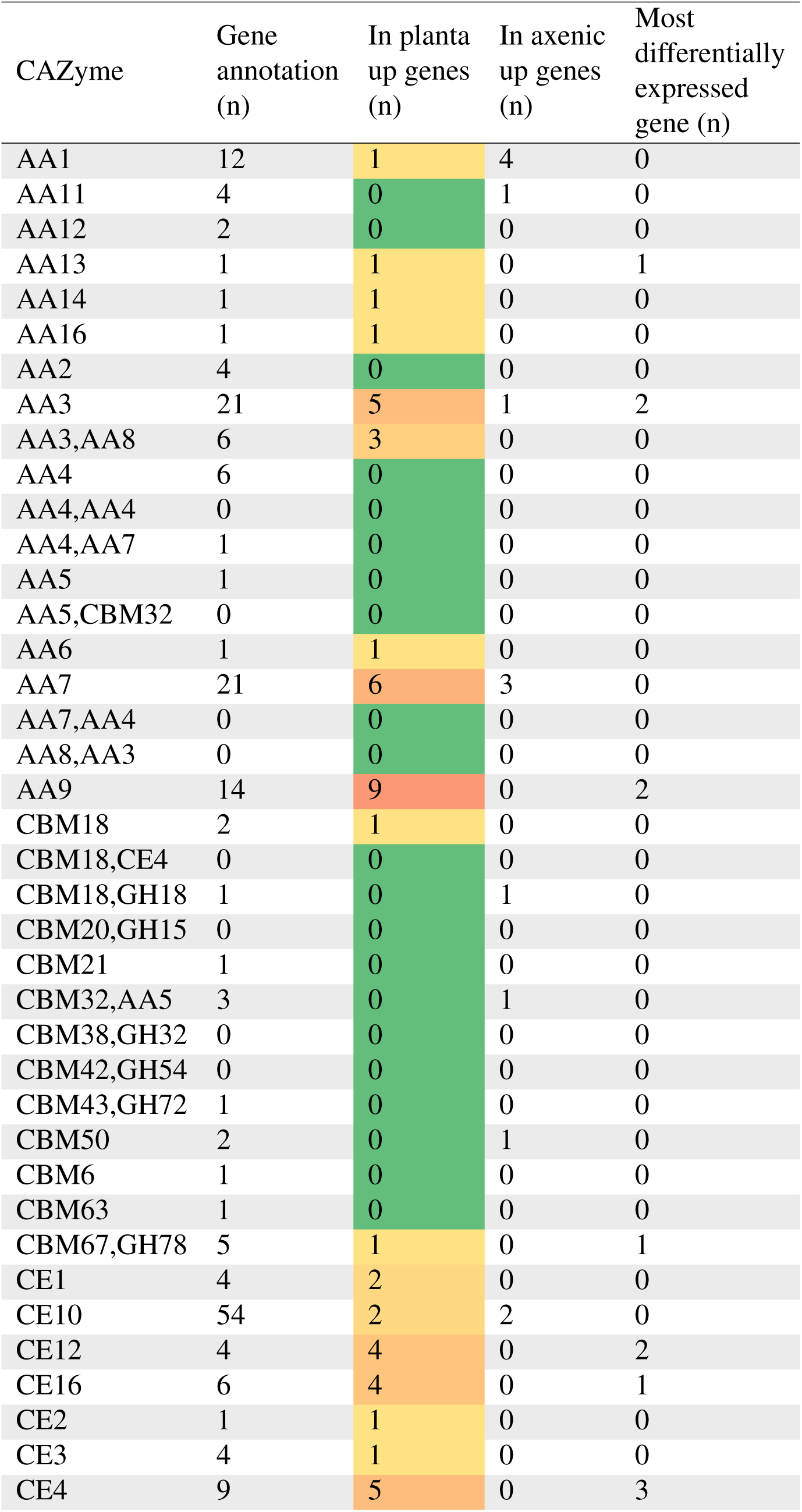
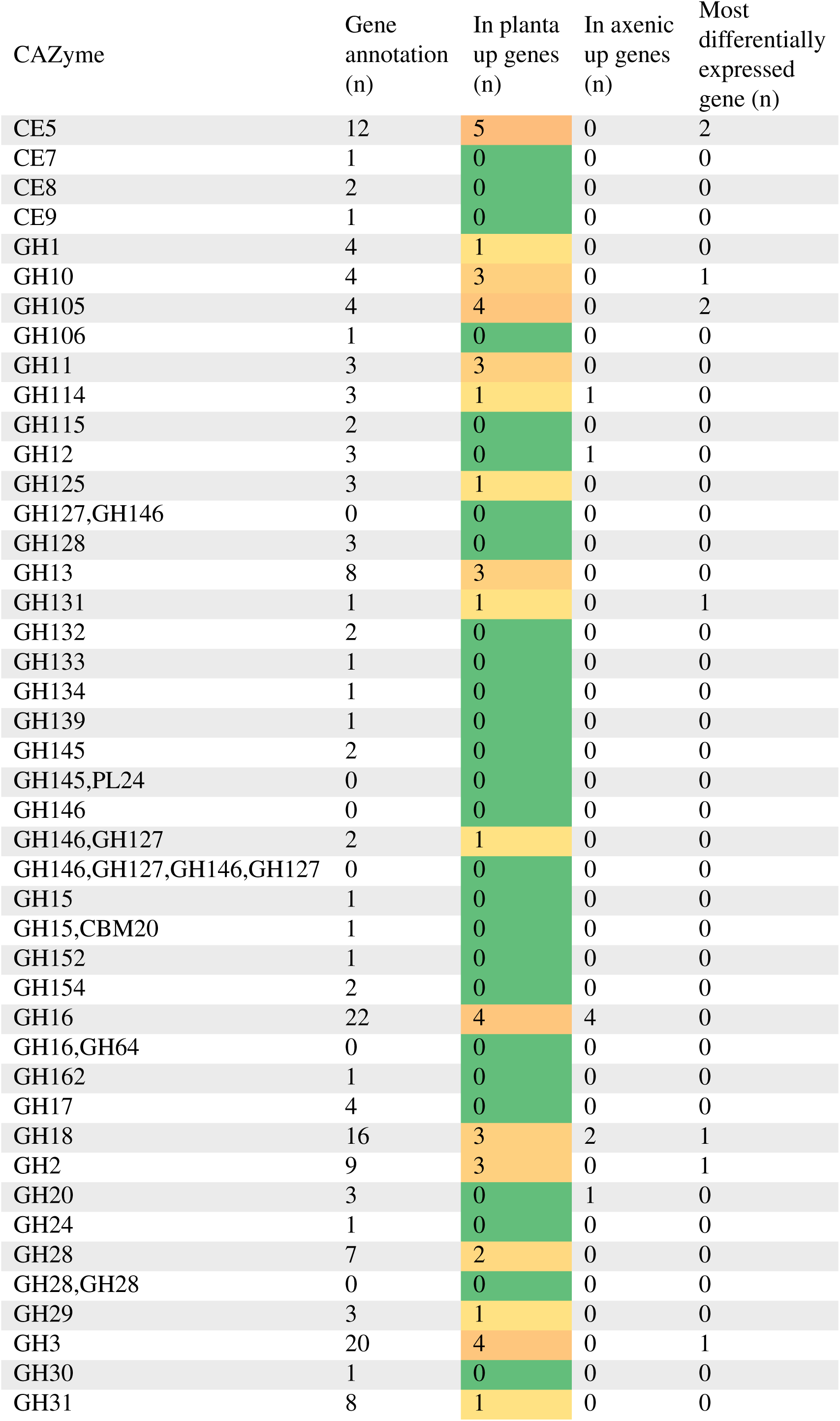
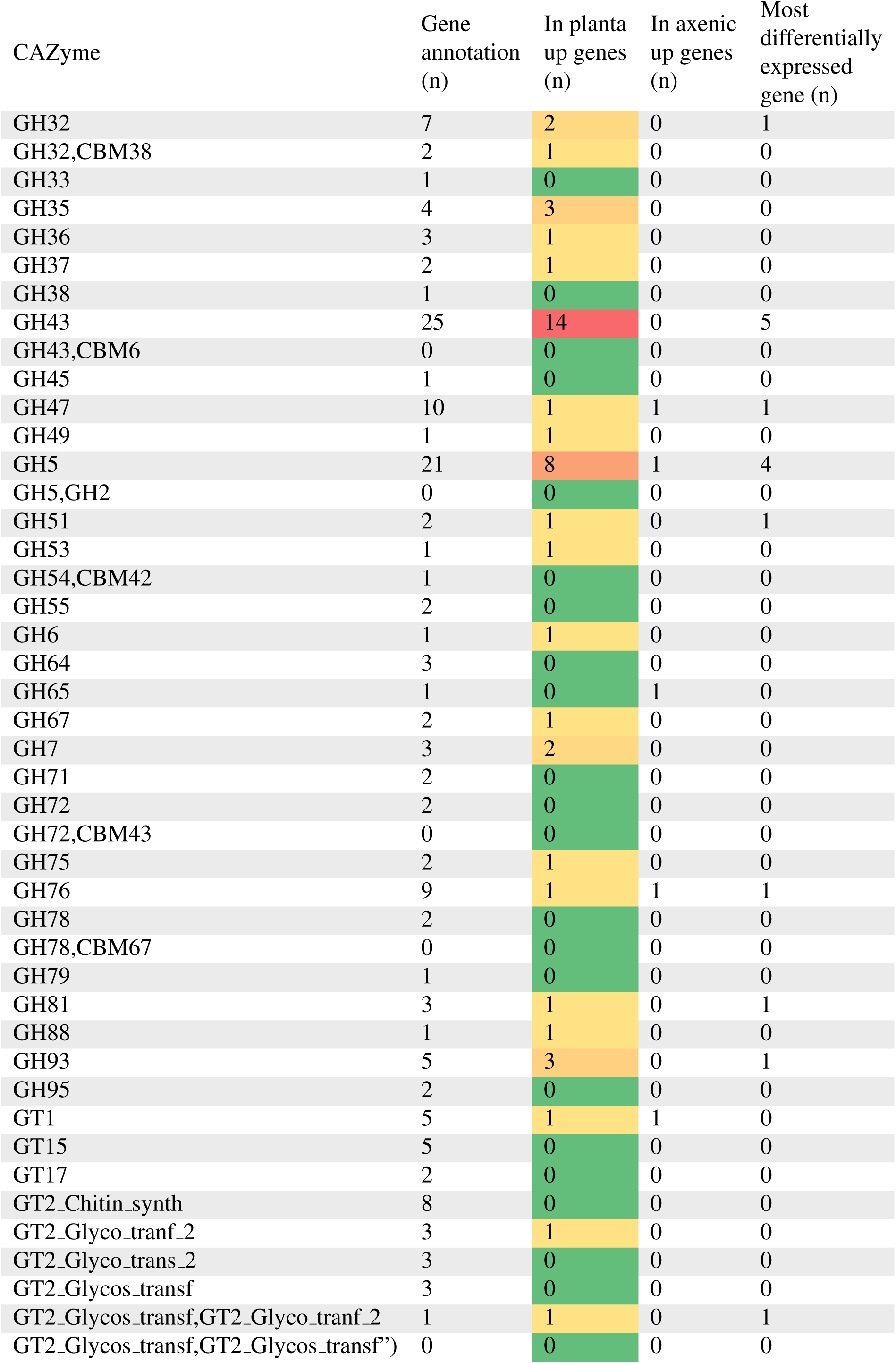
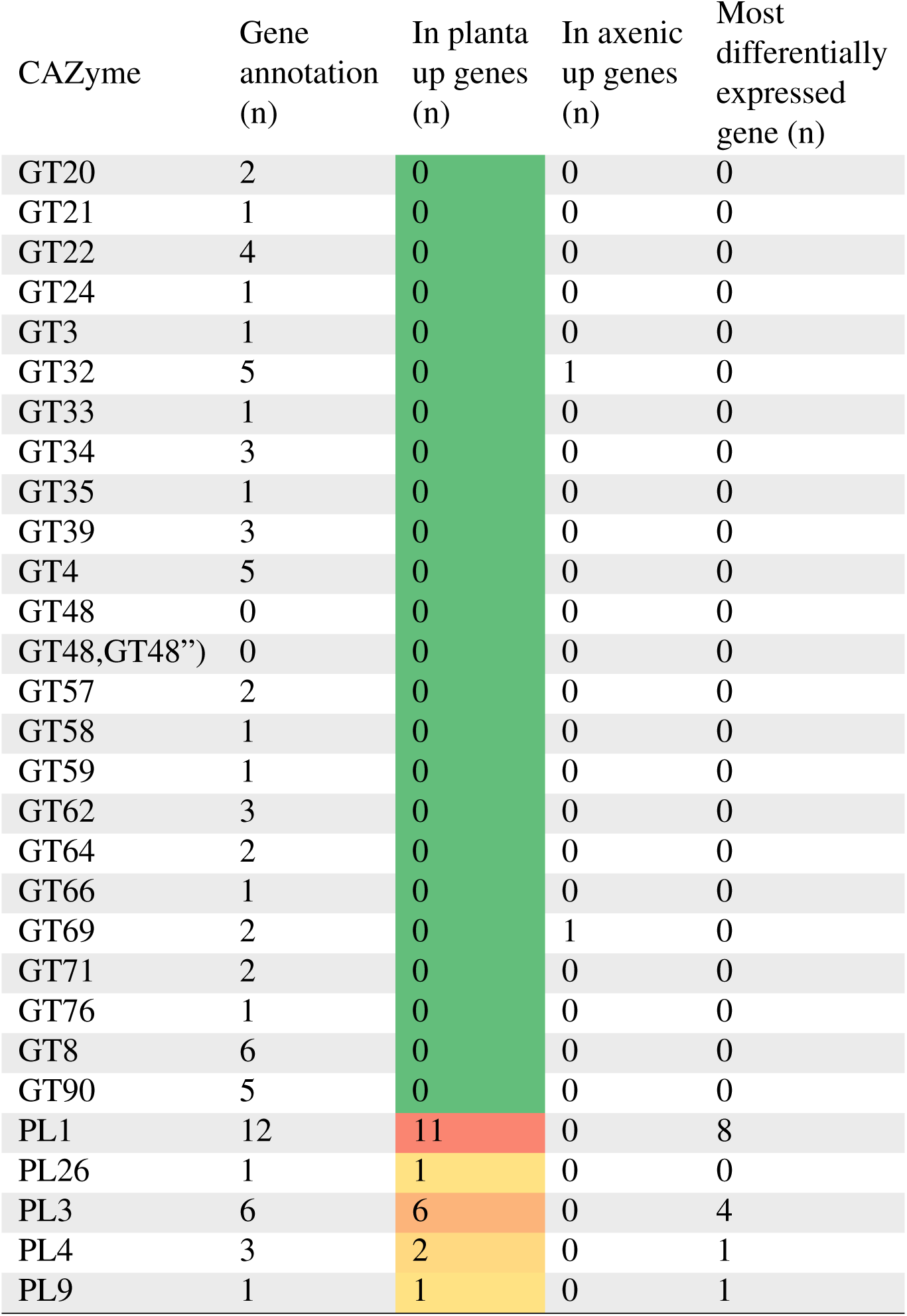
The number of genes expressed for each carbohydrate-active enzyme sub-family in *Fusarium xylarioides* arabica563. The *in planta* up-regulated genes are shaded, where the warmest colours represent the highest up-regulated gene count.

**Table S10:**
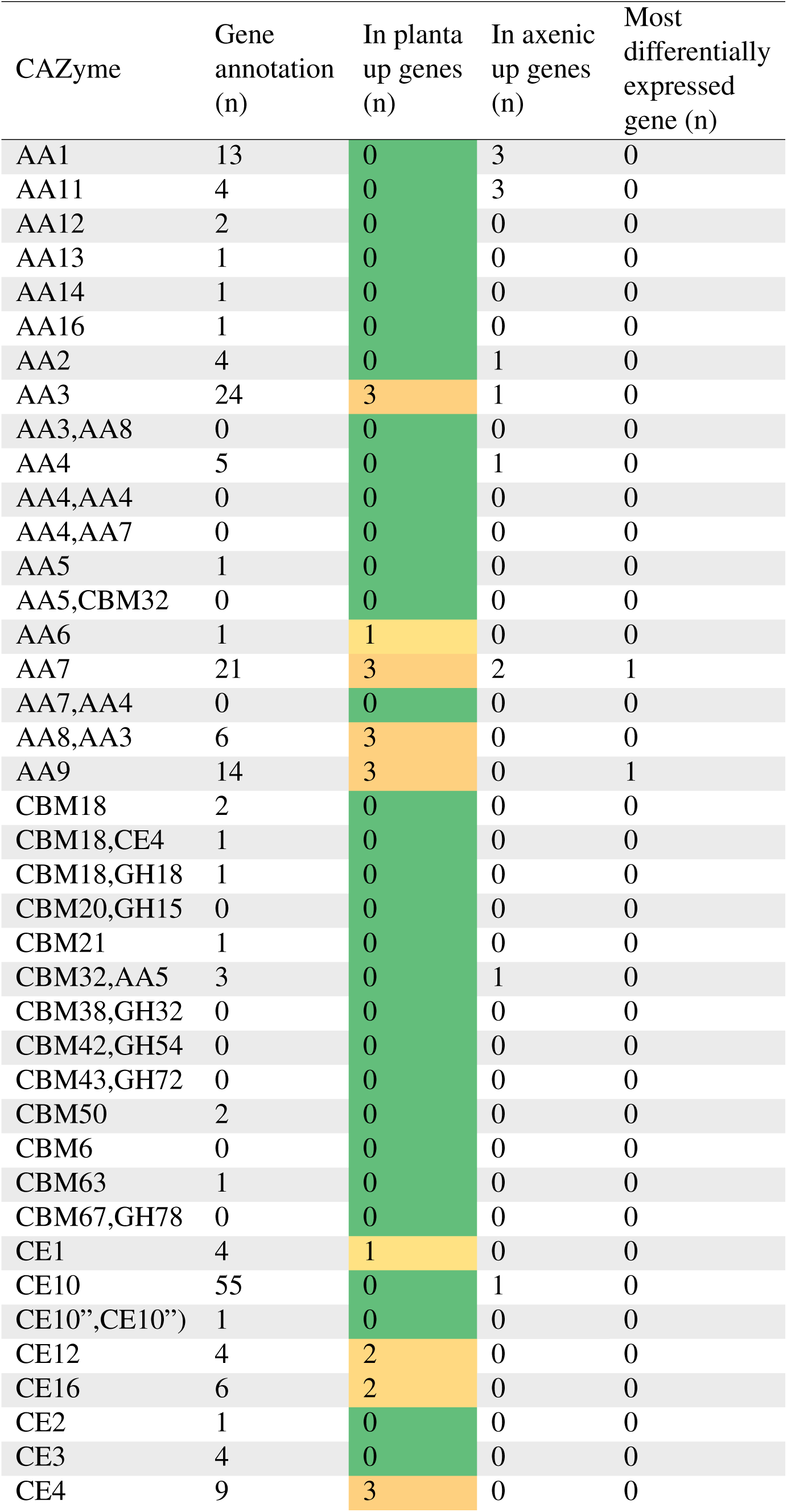
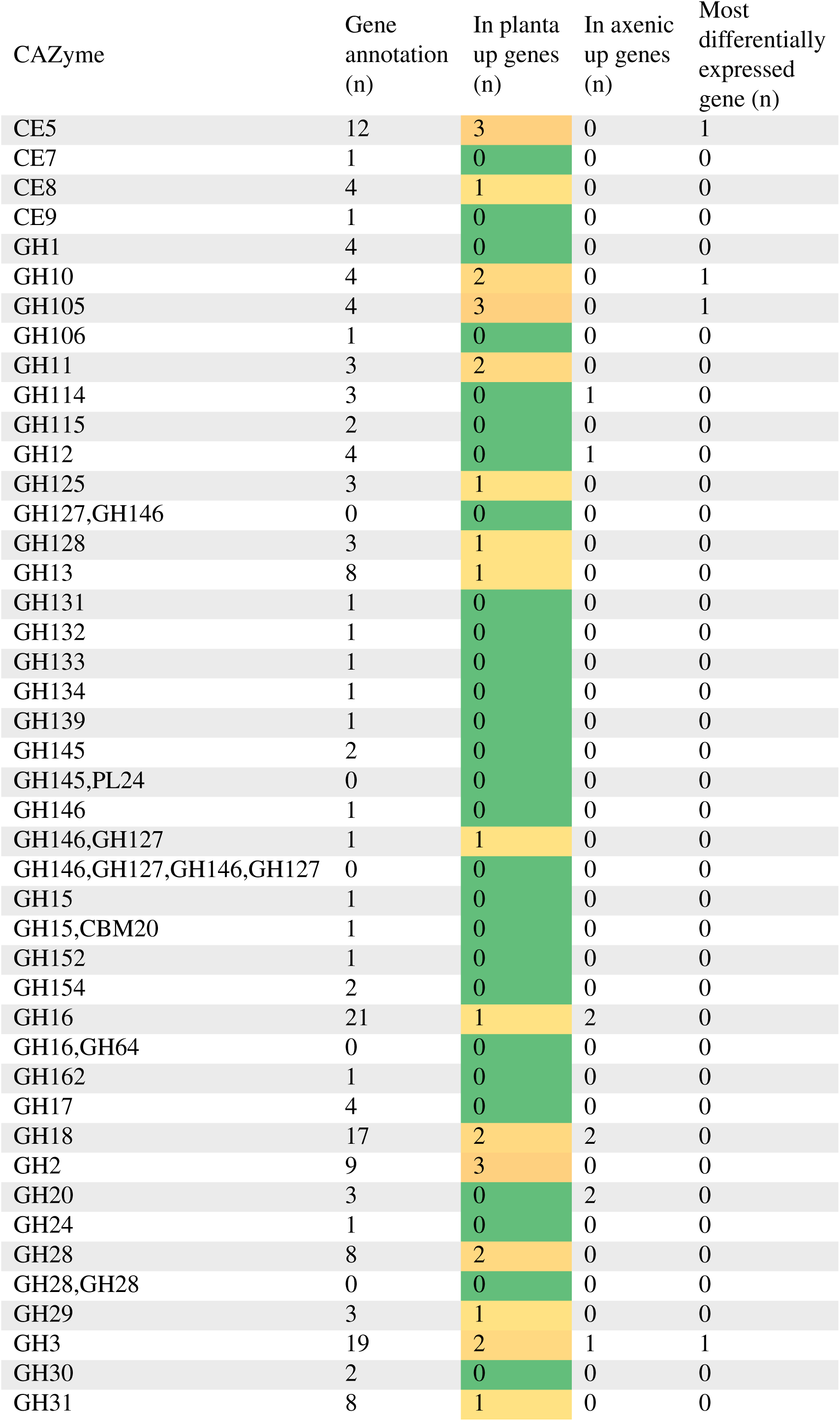
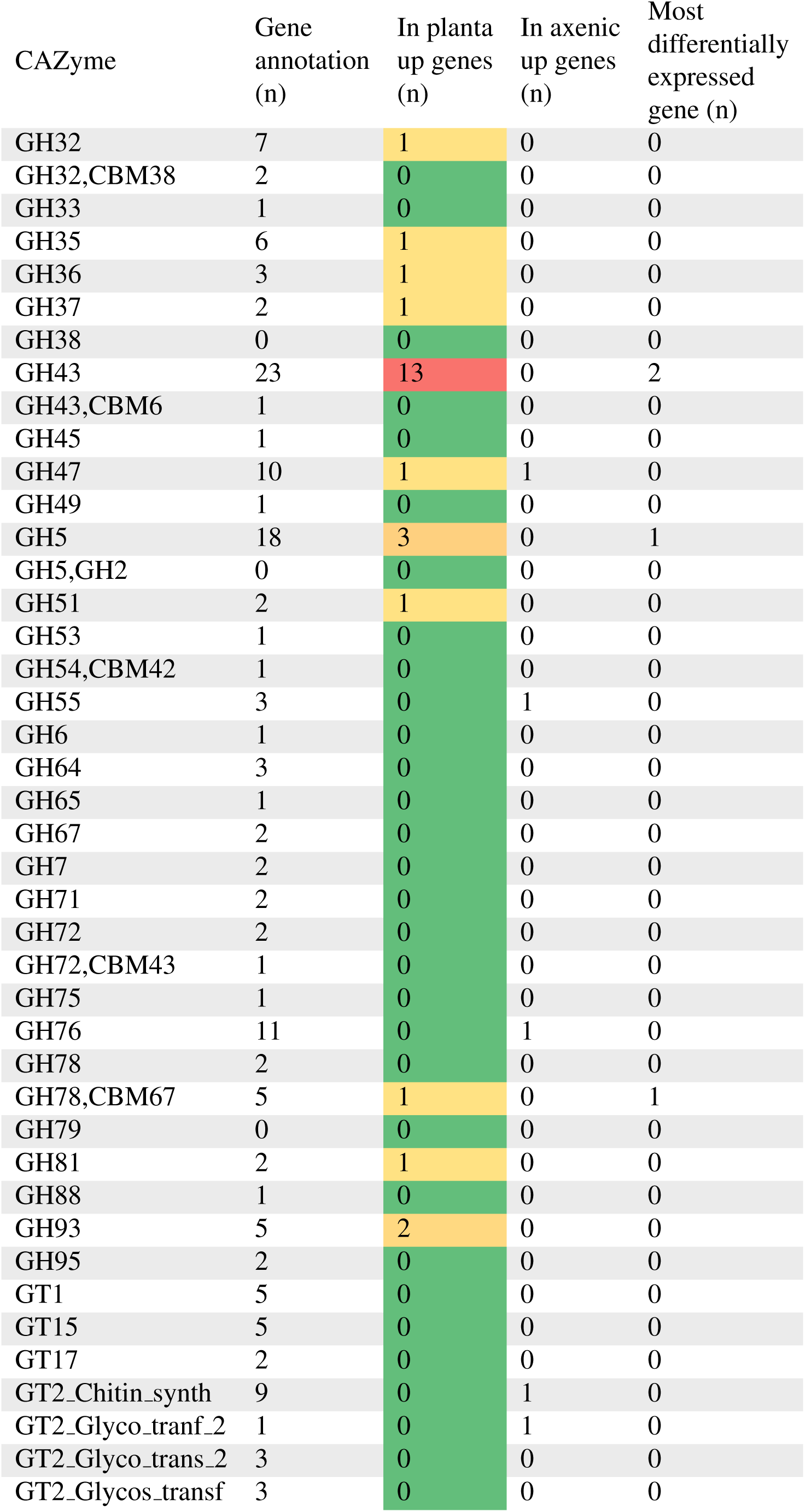
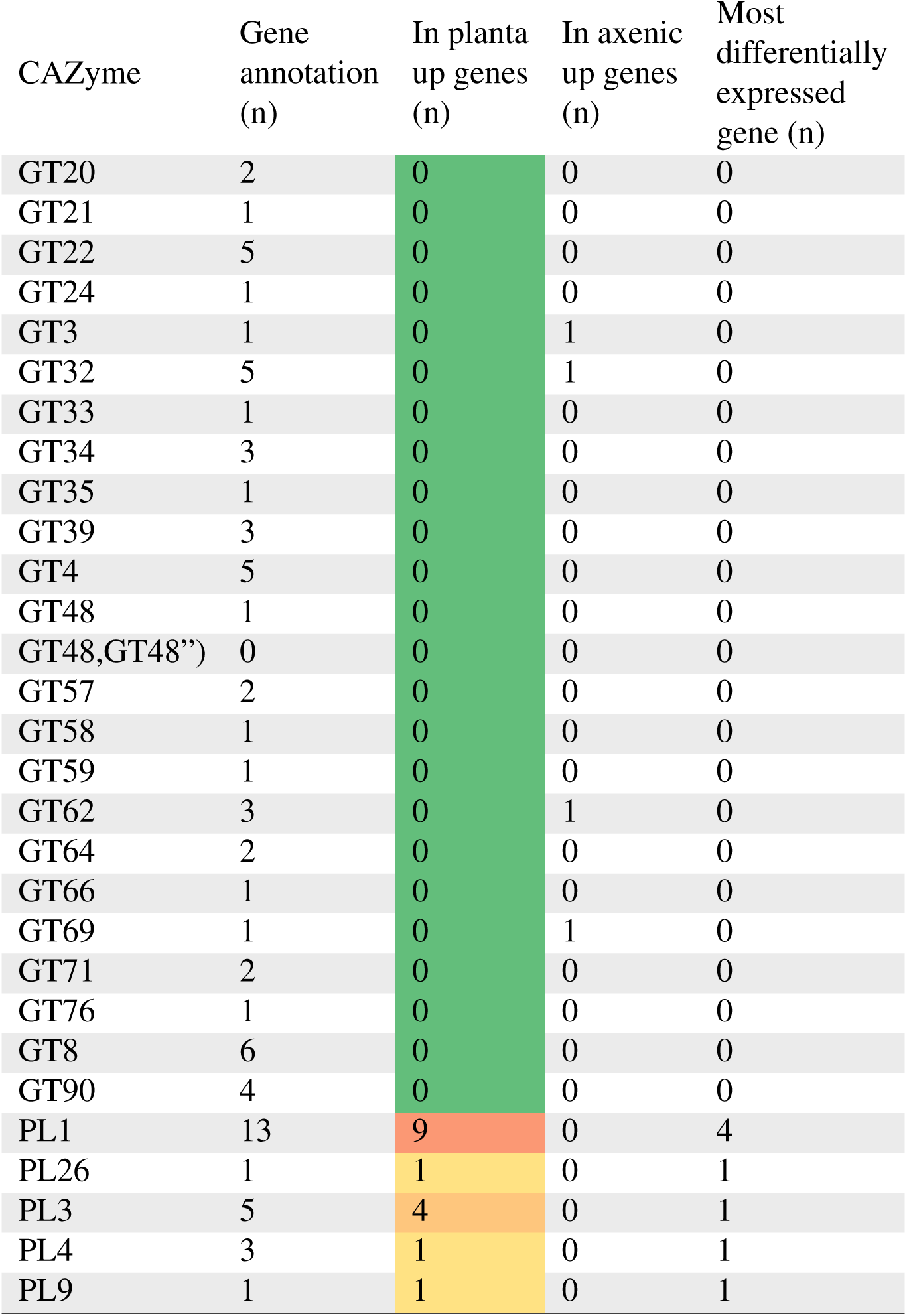
The number of genes expressed for each carbohydrate-active enzyme sub-family in *Fusarium xylarioides* arabica908. The *in planta* up-regulated genes are shaded, where the warmest colours represent the highest up-regulated gene count.

**Table S11:**
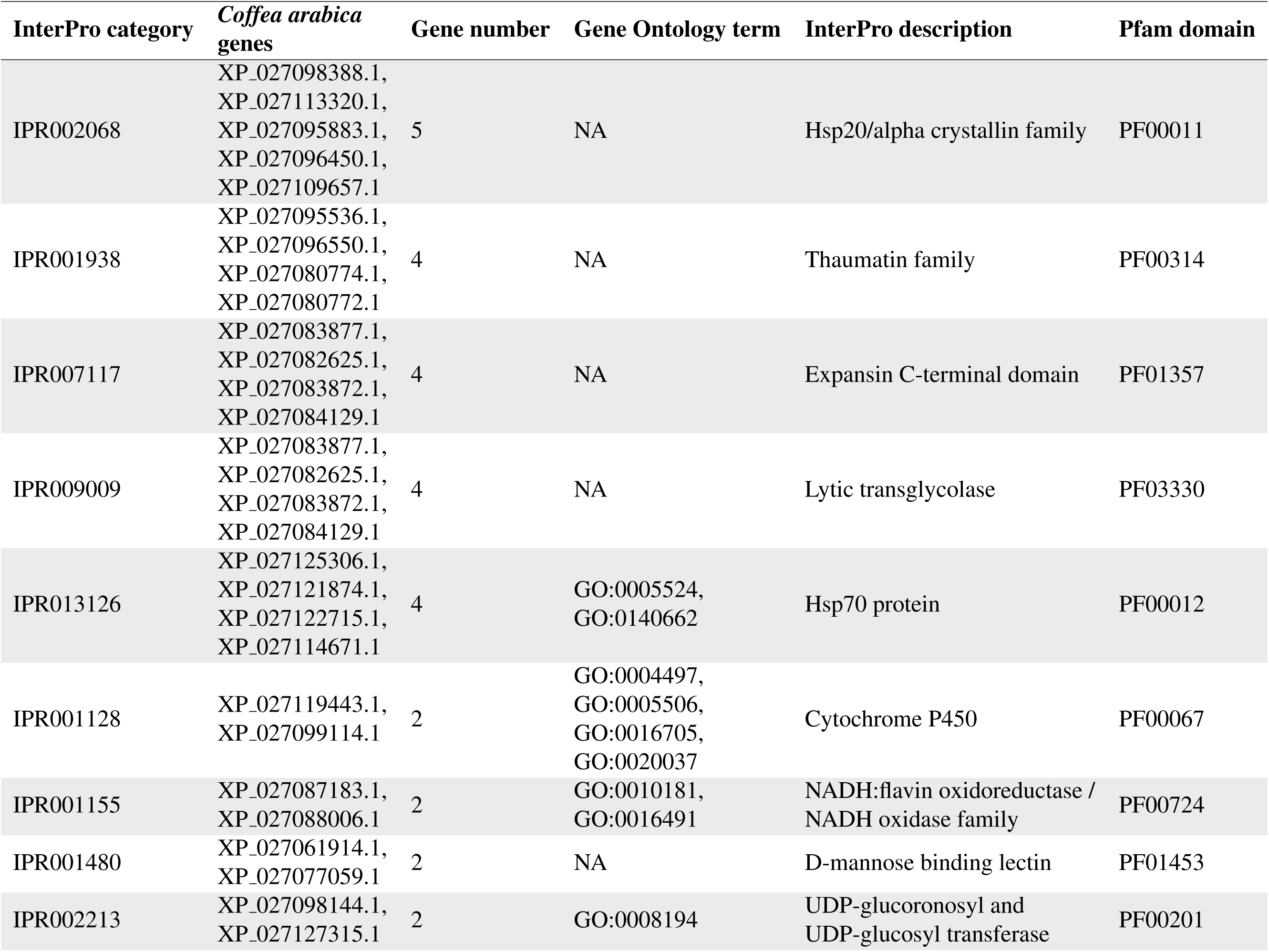
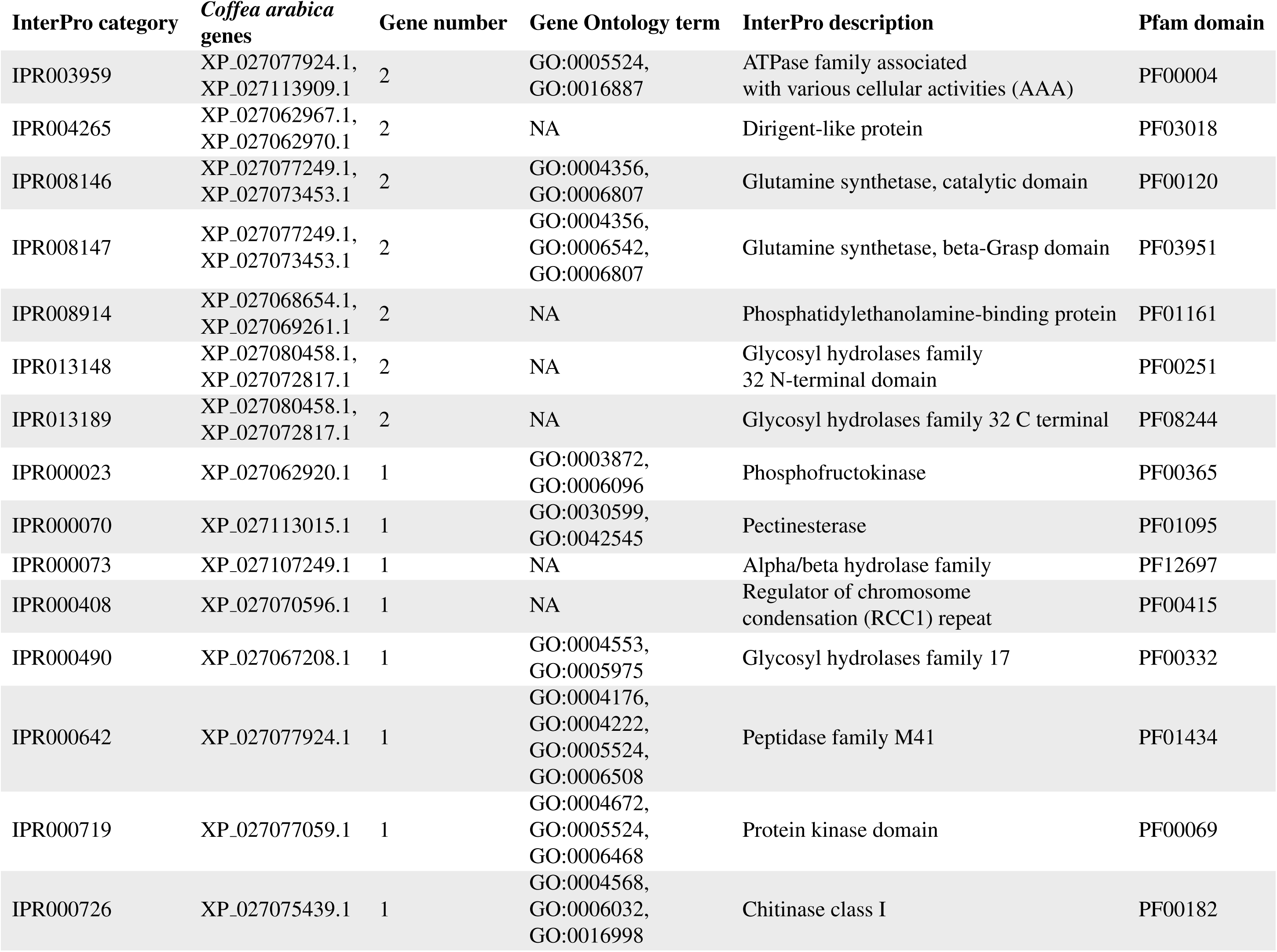
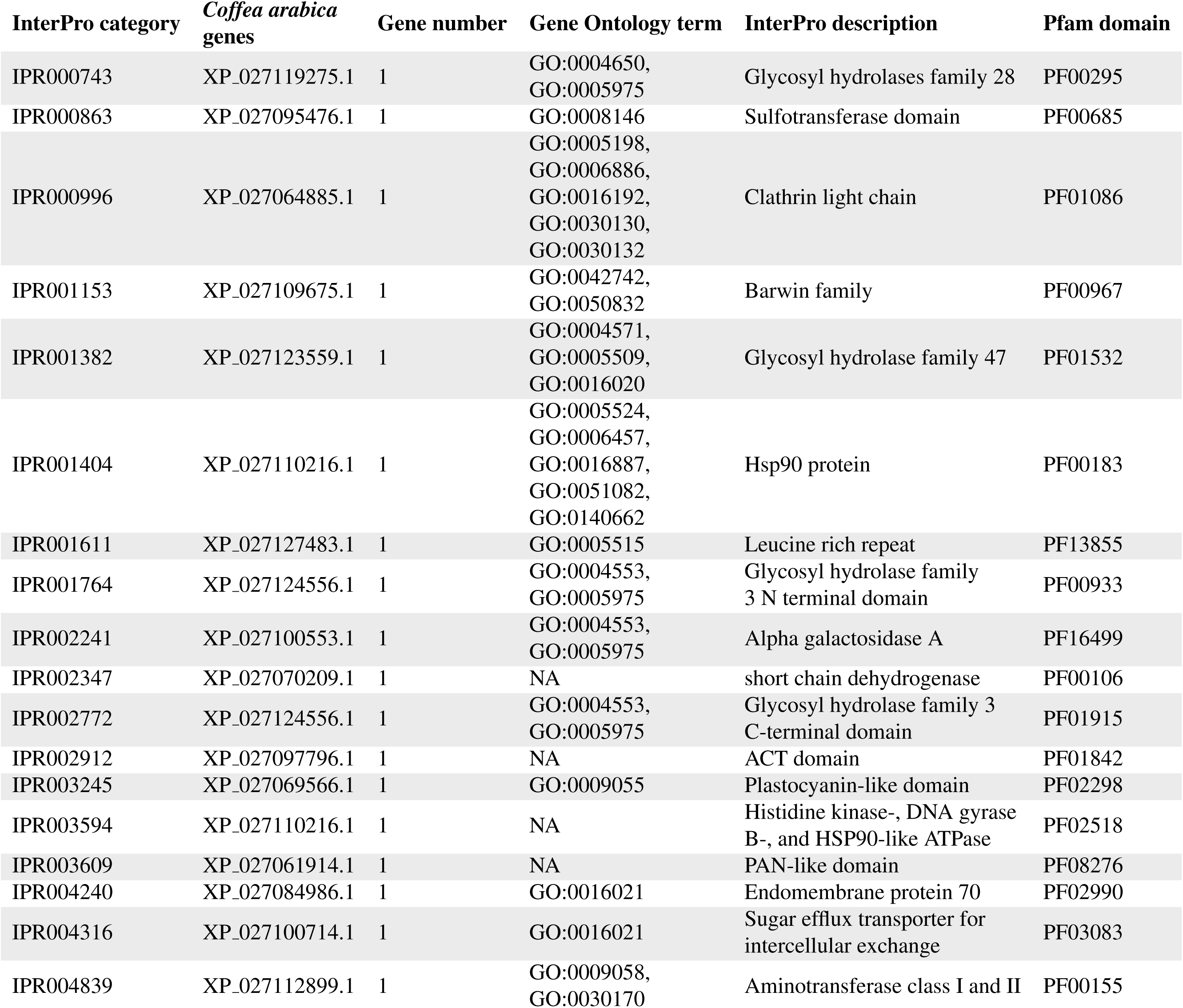
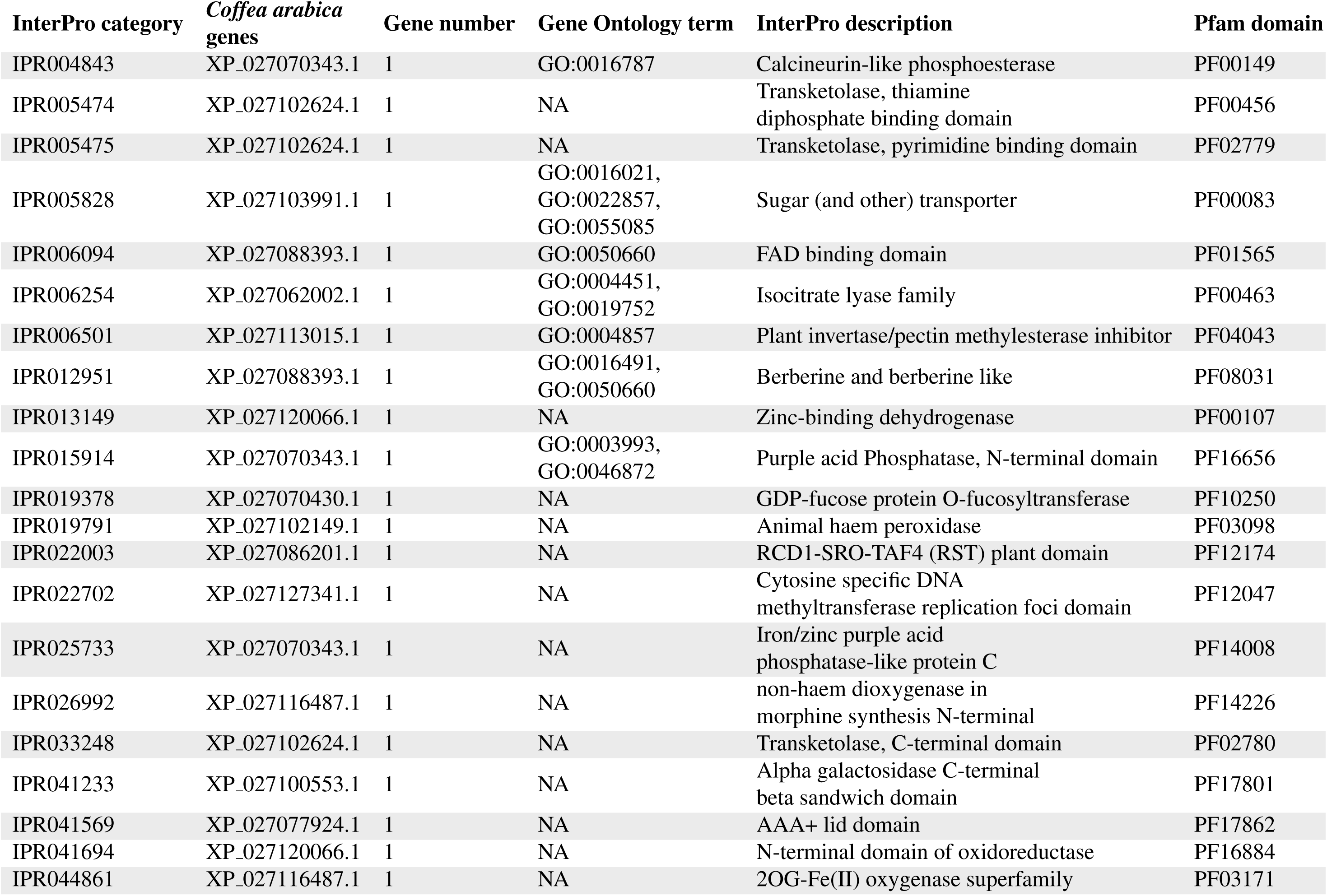
Gene count and description for significantly enriched InterPro terms in the Coffea arabica in planta samples.

**Table S12:** Transposable element density in horizontally transferred regions (HTR), *Starships* and genome-wide in the *Fusarium xylarioides* arabica563 reference. Low complexity and simple repeats were excluded from the transposable element (TE) count.

|  | TE count | Length (bp) | TE/bp |
| --- | --- | --- | --- |
| HTR 1 | 12 | 5,132 | 0.234 |
| HTR 2 | 221 | 240,000 | 0.092 |
| HTR 3 | 15 | 40,000 | 0.038 |
| HTR 4 | 169 | 280,000 | 0.060 |
| HTR 5 | 23 | 24,633 | 0.093 |
| Starship 1 | 71 | 120,000 | 0.059 |
| Starship 2 | 827 | 1,037,016 | 0.080 |
| Starship 3 | 84 | 161,594 | 0.052 |
| Genome | 9 | 100,000 | 0.009 |

